# Omics-driven Identification of Candidate Genes and SNP markers in a Major QTL Controlling Early Heading in Rice

**DOI:** 10.1101/2025.03.25.645207

**Authors:** Deepti Rao, Nikhila Sai T., C.G. Gokulan, Namami Gaur, Mohammed Jamaloddin, Sheshu Madhav Maganti, Meenakshi Sundaram Raman, Hitendra K. Patel, Shrish Tiwari, Ramesh V. Sonti

## Abstract

Precise control of heading date is essential for optimizing regional adaptability, enhancing climate resilience, and maximizing grain yield in rice, making it a key breeding target. The SM93 rice line exhibits a 7-10-day earlier heading than the elite Indian variety, Samba Mahsuri (SM). F2 populations derived from a cross of SM with SM93 were phenotyped for heading date across three successive kharif (rainy) seasons (2019-2021), revealing consistent early heading and notable transgressive segregation. Initial QTL-seq analysis identified a high-confidence QTL on Chr3, *qDTH3*, strongly associated with days to heading (DTH). KASP assays and association mapping were subsequently performed to refine the QTL, narrowing it down to a 2.53 Mb region, explaining ∼25% of the trait. SNP markers closely linked to heading were identified. Transcriptomic analysis revealed significant differential regulation of genes within *qDTH3* and upregulation of the MADS-box network in SM93 during panicle initiation, suggesting their role in promoting early heading. Our integrative strategy led to the identification of candidate genes within *qDTH3*, associated with heading date, phytohormone regulation, and protein turnover, include the transcription factors, *OsMADS34/PAP2* (*Os03g0753100*), *OsZHD11* (*Os03g0718500*), *HSFA2A* (*Os03g0745000*), *OsH3* (*Os03g0727200*), *OSH1/Oskn1* (*Os03g0727000*), a histidine kinase, *HK4* (*Os03g0717700*), and a Serine carboxypeptidase, *OsSCP19* (*Os03g0730400*). Our findings provide SNP markers in and around these genes, as potentially valuable tools for breeding early-maturing rice varieties with improved adaptability.

**HIGHLIGHT:** This study on the SM93 rice line identifies a major QTL for early heading and describes genes and SNP markers that are closely linked to this trait, which can be used in trait advancement. By fine-tuning heading date, we can help rice varieties better withstand temperature extremes, drought, and other climate-related stresses, making it a key target for genetic improvement and breeding strategies.

## INTRODUCTION

Heading date, a crucial trait for crop reproduction, yield, and adaptability, is governed by a complex interaction of environmental cues and internal signals. While *Arabidopsis* research has provided insights into photoperiodic signaling and circadian rhythms, studies in rice, a facultative short-day plant, have revealed several key genetic factors affecting heading date. The regulation of heading date in rice is a multifaceted process driven by a diverse array of genes, including transcription factors, florigens, kinases, phytochromes, and epigenetic factors. Major Quantitative Trait Loci (QTL) such as *Hd1*, *Hd3a*, and *Ghd7* are pivotal in regulating heading date, with *Hd1* and *Hd3a* influencing photoperiod sensitivity and *Ghd7* modulating the plant’s response to light (Shojiro Tamaki et al., 2007; Sun et al., 2022; Yano et al., 2000). The regulation of heading date in rice differs notably under long-day (LD) conditions, being more intricate compared to the simpler mechanisms under short-day (SD) conditions (Han et al., 2016; Komiya et al., 2009). Moreover, the genetic background significantly impacts heading date, as different allelic combinations of genes like *Hd1*, *Ghd7*, *Ghd8*, and *Ghd7.1* can produce varied phenotypic outcomes (Zhang et al., 2019). This complexity arises from the interactions among heading date genes and their dependence on genetic backgrounds, the detailed knowledge of which will enable the development of superior rice varieties that achieve optimal heading dates and enhanced productivity within those same backgrounds.

The rice line SM93 exhibits early heading, leading to maturity approximately ten days earlier than the elite variety Samba Mahsuri. To explore the genetic basis of this trait, we developed F2 populations from a cross between SM93 and Samba Mahsuri and employed the QTL-seq analysis to identify key genomic regions associated with early heading. Additionally, we derived key SNP markers from these QTL regions. Transcriptomic profiling of SM93 revealed differential gene regulation in key pathways related to inflorescence development and hormonal signaling. These findings offer valuable insights for fine-tuning heading date in rice, which is crucial for improving climate resilience and optimizing yields in varying environmental conditions.

## MATERIALS AND METHODS

### Phenotyping of SM and SM93

Inbred SM and SM93 plants were phenotyped in our fields at CSIR-CCMB Annexe-II Hyderabad, India (Latitude: 17.4097, Longitude: 78.5496) for various traits, such as days to heading (DTH), plant height, number of panicles, panicle length, exsertion and branching, and grain number per panicle in Kharif, 2018). DTH was calculated as the number of days from sowing to the emergence of the panicle from the flag leaf sheath. The heights of plants were measured at maturity from the soil till the tip of the flag leaf. To quantify the panicle exsertion trait, the panicle exsertion length variation was measured in SM and SM93 at maturity. Precisely, the panicle exsertion length was measured from the panicle node to the flag leaf collar. A panicle that exerts itself completely has the panicle node exposed towards the outside and the panicle exsertion in this case was expressed as a positive integer. Contrary to this, a choked panicle has its node covered by the flag leaf sheath, in which case, the extent of choking was expressed as a negative integer. The phenotyping data was recorded in Google sheets and MS excel. The mean and standard deviation were calculated and statistical inferences were drawn for all the traits using Welch’s *t*-test at a significance level of *p* <= 0.01.

### DNA Isolation and Sequencing of SM and SM93

Leaf samples from SM and SM93 plants were used for DNA isolation by the CTAB method (Doyle & Hortorium, 1991). Two pools of DNA samples were made; one pool consisting of DNA from the leaves of 6 SM plants, and another pool containing DNA from the leaves of 6 SM93 plants. Paired-end short read sequencing of the DNA samples was outsourced to Nucleome Bioinformatics Pvt. Ltd., Hyderabad, wherein, whole genome sequencing was performed on the Illumina HiSeq2500 platform.

### Variant Discovery in SM93

FASTQC (v0.11.2) was used to assess the quality of raw NGS reads. The reads were trimmed with a threshold of Q20 (BBDuk2 v37.36). After trimming, the reads shorter than 20 bases were filtered out to reduce ambiguous mapping to the reference (BBDuk2 v37.36). The resulting high-quality reads were ready to be used for variant analysis. The high-quality SM and SM93 reads were mapped individually to the IRGSP-1.0 genome assembly using BWA (v0.7.17), after which the SAM output was converted to BAM format using Samtools (v1.3.1). Then, the bam files were processed to exclude PCR duplicates and the resulting files were indexed with Samtools (v1.3.1). Variants were called using FreeBayes (v1.1.0) in the form of a VCF file. The variants were then filtered using a depth threshold of 10 and mapping quality threshold of 40, using a Perl script. InDels were filtered for RO=0, LEN=10 and GT=1/1. The high-quality variants thus obtained were annotated using Variant Effect Predictor (VEP) and visualized with Integrated Genomics Viewer (IGV). Primer sets were designed to amplify regions of insertions and deletions discovered by NGS analysis for F1 hybridity testing.

### Cross-pollination of SM and SM93

Healthy plants from the field were transplanted to pots after the panicles had emerged. The pots were maintained in our greenhouse at 30° C, and crosses were performed with SM as the female parent and SM93 as the male parent. Putative F1 (hybrid) seeds resulting from the crosses were collected, germinated and raised in the greenhouse, alongside the parents, SM and SM93. For the initial 15 days, we transplanted the plantlets to artificial soil, comprising soilrite, perlite and vermiculite (1:1:1). The artificial soil was supplemented with half MS, once in a week. After 15 days, the plants were moved to larger pots with clayey soil and retained in the same pots until maturity. Leaf samples were collected from 1-month old plants for DNA isolation by the CTAB method (Doyle & Hortorium, 1991). Hybridity of the putative F1 plants was tested using the insertion and deletion markers developed based on InDel data. F1 plants confirmed for hybridity were grown up to maturity. We recorded DTH data for the F1 plants and their parental controls. Natural, self-pollination resulted in F2 seeds, which were collected from each F1 hybrid in separate paper covers and maintained in the seed storage room (4℃) for further use.

### F1, F2 and F3 Phenotyping and F2 Extreme Bulk Sampling

F2 seeds from one paper cover (single F1 plant) each were grown in the field and phenotyped for DTH in Kharif 2019, 2020 and 2021, along with their parental controls. For each F2 plant, DTH was recorded only once, at the emergence of its first panicle. In the three years of our study, 130, 131 and 121 plants were phenotyped, and the DTH frequency was plotted every time. The plants lying in the tails of the F2 distribution in the plot were selected for QTL-seq analysis. To be specific, the plants lying in the left tail of the distribution comprised the Early Bulk (EB), while those lying in the right tail of the distribution made up the Late Bulk (LB).

Additionally, the F3 seeds of the EB and LB F2 plants were sowed from the Kharif 2020 population. The progeny of each F2 EB and LB plant was grown in smaller separate plots in the same field to understand if the DTH trait was stable in the next generation (F3). Statistical significance testing for heading date variation was performed using the Welch’s *t*-test. Plants from the early bulk are being tested in advanced generations for trait stability.

### Short-read Sequencing for QTL-seq

Equal weights of leaf tissues (stored previously at -80 °C) of EB plants were ground together using liquid nitrogen and processed as per Doyle & Hortorium, 1991 to isolate EB DNA. Similarly, the DNA sample for the LB was prepared from the LB plants. 8 plants from both extremes were taken for early and late bulks in Kharif 2019. 12 and 13 plants were taken as early and late bulk, respectively, in Kharif 2020, while 11 plants from both the extreme bulks were taken in Kharif 2021. Whole-genome sequencing of all the EB and LB DNA samples was performed on Illumina NovaSeq 6000 at 40X coverage to obtain population-specific, bulk-wise short-read data for QTL-seq analysis.

### QTL-seq Analysis

The Takagi et al. QTL-seq pipeline was followed to identify QTLs in SM93 (Takagi et al., 2013). Short-reads of the Kharif 2019 EB and LB, and parent cultivars were mapped to the IRGSP-1.0 (Nipponbare) reference genome with BWA (v0.7.17). BAM files obtained were sorted, duplicates were removed and the resulting BAM files were indexed with Samtools (v1.3.1). The qtlseq and qtlplot scripts of the QTL-seq (v2.2.3) pipeline (Sugihara et al., 2022) were then run by opting for 10,000 iterations to obtain the 95% and 99% confidence intervals (CIs). Statistical CIs of Δ SNP-index for all the SNP positions with given read depths under the null hypothesis of no QTLs were calculated. QTL identification was based on the resulting plot and the **Δ** SNP-index values corresponding to the peaks or troughs obtained at p < 0.05 under the null hypothesis. The same method was followed to detect QTLs based on the Kharif 2020 and 2021 EB and LB short-reads. In addition, G analyses (Mansfeld & Grumet, 2018) were performed on the data from the three populations. Subsequently, we also repeated the QTL-seq analyses using the SM reference genome.

### Validation of SNPs and Association Mapping

Kompetitive Allele Specific PCR (KASP^TM^) assays (LGC Genomics, UK) were used for validation of SNPs and association mapping. Common SNPs among the 3 mapping populations in *qDTH3* region were identified. To refine our dataset, we selected high-confidence SNPs based on SNP annotation and Δ SNP values, most of the time, prioritizing those with Δ SNP values greater than 0.7 across all the three populations. Additionally, we included some SNPs located in known heading date/flowering QTLs based on QTARO (QTL Annotation Rice Online database) data available on SNP-seek (https://snp-seek.irri.org/) database, known developmental and flowering-related genes from our literature survey, as well as those with impactful consequences, such as missense mutations, start lost variants, and 5’ and 3’ UTR variants. SNPs predicted to be deleterious by SIFT were also given priority in our analysis. Some SNPs were selected as fillers between distant SNPs to enhance the resolution of our mapping efforts. A total of 49 SNPs, which comprised 47 (KASP^TM^ test SNPs) SNPs from *qDTH3*, and 1 SNP each from another region of Chr3 and Chr1 (KASP^TM^ negative control SNPs) were finally selected for KASP^TM^ assays. Genotyping was performed in a 5 µl reaction per assay under conditions recommended by the manufacturer. KASP^TM^ reaction condition used was as follows: 94 ℃ - 15 minutes; 10 cycles of 94℃ - 20 seconds and 61-55℃ - 1 minute (touch- down - 0.6℃/cycle); 26 cycles of 94℃ - 20 seconds and 55℃ - 1 minute in the ABI ViiA7 RT-PCR System or Bio-Rad CFX384 system. SNP markers were initially screened in SM and SM93 to identify nucleotide variations between the two lines. Selected markers were then tested in EB and LB plants from the Kharif 2019 population to assess their consistency with the phenotype. Finally, all F2 individuals were genotyped for the final test SNPs and control SNPs. Association mapping was conducted using KASP SNP markers and phenotypic data for heading date in TASSEL 5.2.88. A mixed linear model (MLM) was used to account for population structure and kinship effects. We noted the *p*-values and *R*^2^ values for each SNP to assess the strength of the association with the trait. Following TASSEL-based association mapping, cluster analysis was performed on the significant SNPs using ChiPlot (www.chiplot.online). A Euclidean distance matrix was generated based on the selected SNPs, and hierarchical clustering was performed using the average linkage method (UPGMA). The resulting dendrogram was visualized in ChiPlot to assess genetic relationships among the genotypes.

### RNA Isolation and Sequencing

In Kharif 2020, the flag leaf sheaths were collected from multiple SM and SM93 plants growing in the field at the panicle initiation stage. Undifferentiated reproductive shoot apical meristem tissues measuring 2-3 mm were carefully isolated by dissecting the flag leaf sheaths longitudinally. Pooled reproductive shoot apical meristem tissue samples of different SM and SM93 plants were stored (at -80℃) as triplicates and subsequently processed. RNA was isolated from the samples using RNeasy Plant RNA Isolation Kit (Qiagen, Germany) with on-column DNA digestion using RNase-Free DNase Set (Qiagen, Germany). Purified RNA was quantified using Qubit^TM^ RNA HS Assay Kit (Thermo Fisher Scientific, USA). RNA integrity was assessed using 1% agarose gel electrophoresis and BioAnalyzer 2100 (Agilent Technologies, USA). 1 µg of total RNA samples was used for library preparation for each sample using the Illumina TruSeq Stranded Total RNA with Ribo-Zero Plant (Illumina, USA). Sequencing was performed on Illumina’s NovaSeq6000 platform with cycling conditions to obtain ∼100 million reads per sample in the form of paired-end (2 x 100 bp) reads.

### Differential Gene Expression Analysis

The quality of the raw data was assessed using FastQC. Adapters and low-quality reads were trimmed using Trim Galore! (v0.6.7). RNA-STAR was used to align the trimmed reads to the IRGSP-1.0 (Nipponbare) rice genome. The reads mapped to genes were counted using featureCounts. Differential gene expression (DGE) analysis between the triplicates of SM93 vs SM was performed using the R program DEseq2. Genes with a false discovery rate ≤ 0.05 and log2 Fold Change (log2FC) ≤ -1 or ≥ 1 were considered high- confidence downregulated or upregulated genes, respectively. Gene ontology analyses of the Differentially Expressed Genes (DEGs) were performed online using RiceFREND (https://ricefrend.dna.affrc.go.jp/), gProfiler (https://biit.cs.ut.ee/gprofiler/gost) and AgriGO (http://systemsbiology.cau.edu.cn/agriGOv2/).

## DATA AVAILABILITY

The raw Whole Genome Sequencing (WGS) data for the parent and bulk samples, as well as the raw RNA-seq data for the parental lines, have been deposited in the NCBI Sequence Read Archive (SRA) under the accession number <u>PRJNA1127325</u>. These datasets are publicly available and can be accessed via the provided link.

## RESULTS

### Phenotyping in Natural Field Conditions Showcased Enhanced Agronomic Traits in SM93

SM93 plants were phenotyped in Kharif 2018 for development and yield traits. The heading date of SM93 was ∼10 days earlier than that of SM (mean values for DAS were 100.1 and 90.25 DAS for SM and SM93, respectively; *p*<0.01, Welch’s *t*-test), leading to earlier flowering and grain filling in SM93 **(*Figure 1A-C*, *Figure 2A, Supplementary data S1)***. Notably, SM panicles partially choked, while SM93 panicles fully emerged and were significantly longer **(*Figure 1E*, *Figure 2D, E, Supplementary data S1)***. There were no significant differences in height, panicle number and primary panicle branching between SM93 and SM ***(Figure 2B, C, and F, Supplementary data S1)***. SM93 also had more secondary branches and grains per panicle **(*Figure 1D*, *Figure 2G, H, Supplementary data S1*)**. From our field observations, we could conclude that SM93 exhibits an earlier flowering pattern compared to SM, which could potentially lead to an early start in grain filling, thereby shortening its overall duration **(*Figure 1 F-H*)**. Additionally, the higher yields of SM93 compared to SM could be due to multiple advantageous panicle traits of SM93.

**Figure 1:**
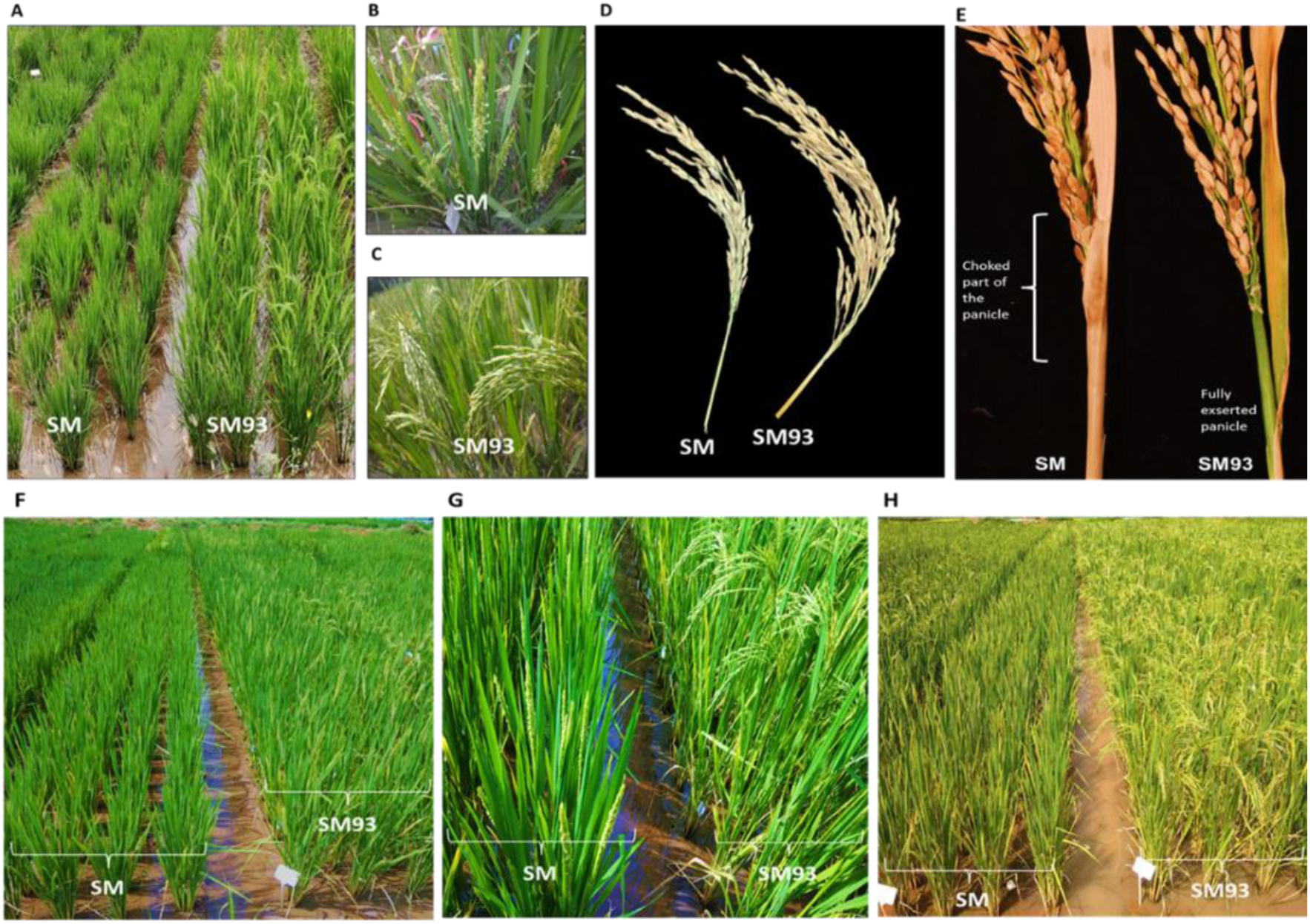
SM vs SM93 in Natural Field Conditions. **(A)** SM in the booting stage and SM93 in the flowering stage at 95 days after sowing (DAS). **(B)** SM at the flowering stage at 103 DAS. **(C)** SM93 at the grain-filling stage at 103 DAS. **(D)** SM exhibits shorter and less branched panicle and SM93 exhibits longer, more branched panicle. **(E)** Incomplete panicle exsertion in SM and complete panicle exsertion in SM93. **(F), (G), (H)** SM and SM93 plant development from the early reproductive stage until grain hardening. **(F)** SM plants in the booting stage and flowered SM93 plants at 95 DAS. **(G)** A closer view of flowered SM and grain-filled SM93 plants at 103 DAS. **(H)** SM plants in the early stages of grain-filling, while SM93 plants are in the grain hardening stage at 120 DAS.

**Figure 2:**
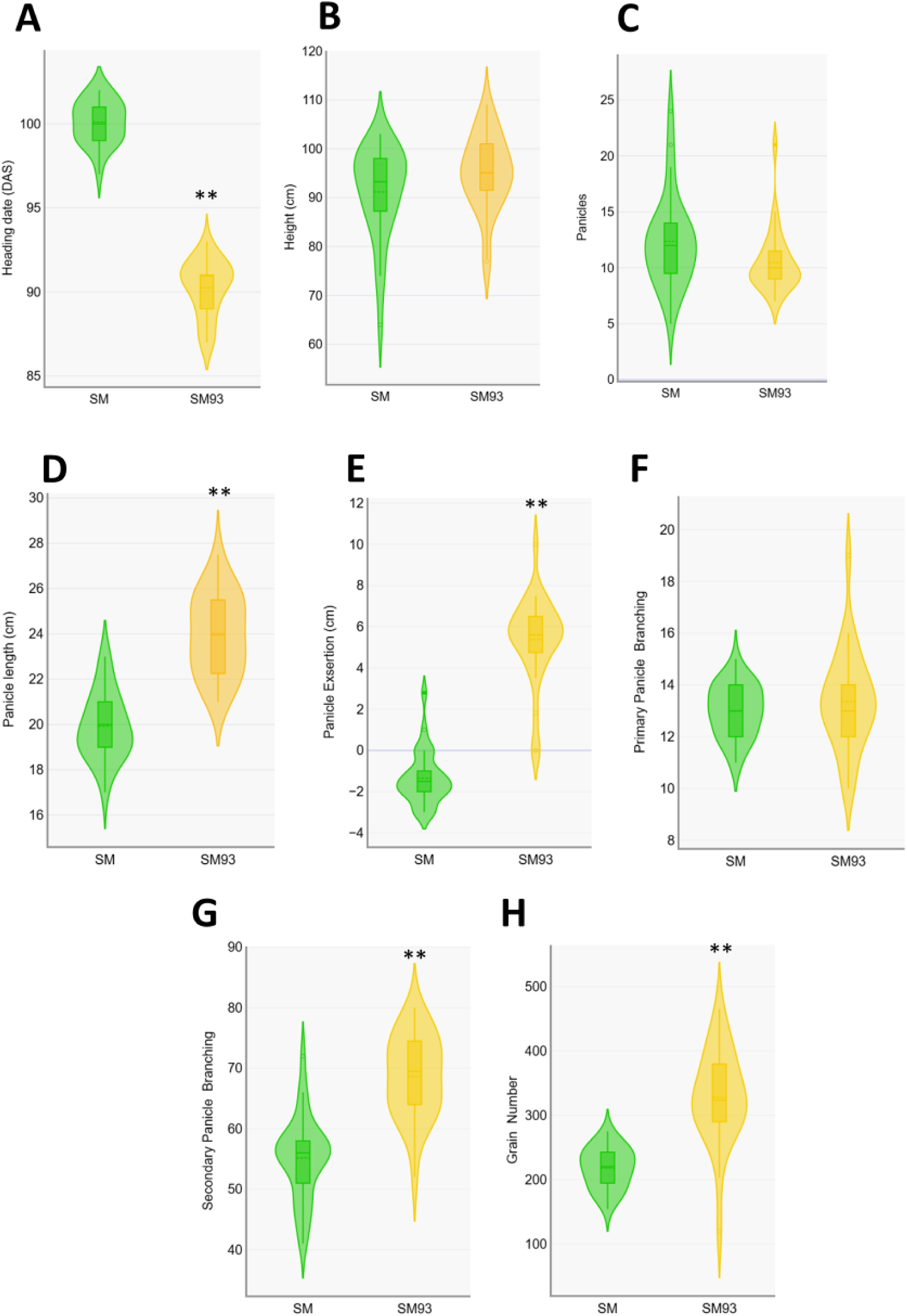
Phenotyping of SM93 for Traits Related to Development and Yield. (A) Days to Heading (DTH) **(B)** Plant height **(C)** Number of panicles per plant **(D)** Length of the panicles **(E)** Panicle exsertion **(F)** Number of primary branches in the panicles **(G)** Number of secondary branches in a panicle **(H)** Grains per panicle. ** indicate p-values < 0.01 using Welch’s T-test (Two-tailed, independent) and n=28.t height **(C)** Number of panicles per plant **(D)** Length of the panicles **(E)** Panicle exsertion **(F)** Number of primary branches in the panicles **(G)** Number of secondary branches in a panicle **(H)** Grains per panicle. ** indicate p-values < 0.01 using Welch’s t-test (Two-tailed, independent) and n=28.

### Segregating Populations Exhibited a Broad Spectrum of Heading Date Phenotypes

DTH was evaluated across multiple generations of SM, SM93, and their progeny in the field. Crosses between SM and SM93, with SM as the female parent, produced F1 hybrids with significantly early DTH values (μ = 100.7 DAS) than both parental lines, SM (μ = 118 DAS) and SM93 (μ = 111 DAS) ***(Supplementary data S2)***. Segregating F2 populations were generated from individual F1 plants, forming the basis for further phenotypic analyses and QTL-seq investigations.

Over three growing seasons (Kharif 2019, 2020, and 2021), 130, 121, and 131 F2 plants were phenotyped alongside SM and SM93 parental controls. Panicles of SM93 consistently headed out earlier than SM, demonstrating phenotypic stability across years ***(Figure S1)***. In Kharif 2019, the DTH values for SM and SM93 were 98 and 90 DAS, respectively. This difference persisted in later years, though heading was slightly delayed: 108 DAS (SM) vs. 101 DAS (SM93) in 2020, and 2021 ***(Supplementary data S3)***. The F2 populations exhibited a wide range of heading dates, revealing continuous variation and transgressive segregation. Heading dates ranged from 81 to 119 DAS in 2019, 86 to 130 DAS in 2020, and 89 to 133 DAS in 2021. A significant proportion of the F2 plants exhibited extreme phenotypes, with 65%, 71%, and 58% of individuals classified as transgressive segregants in the three years, many plants exhibiting earlier heading than SM93 **(*Figure 3, Supplementary data S3)***.

**Figure 3:**
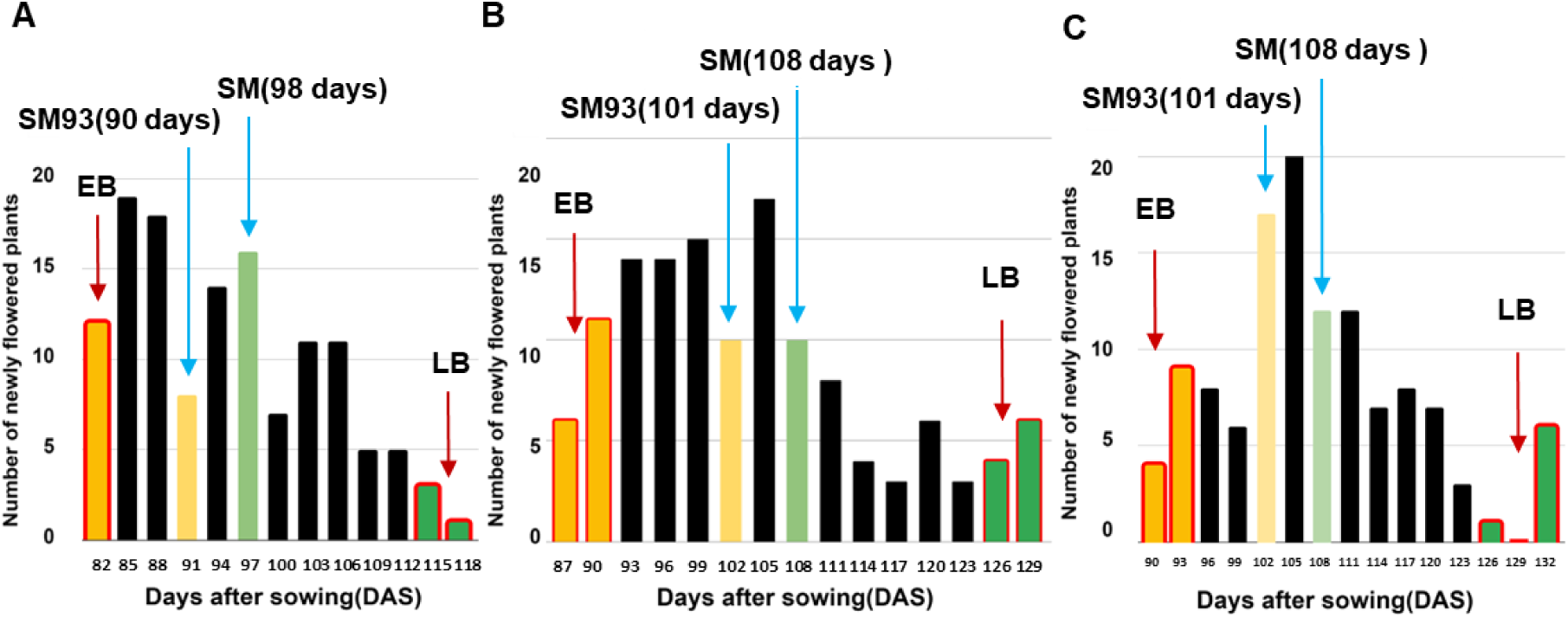
DTH Distribution of the F2 Population (SM x SM93). (A), (B) and. **(C)** show the DTH distribution in the 3 F2 populations screened in the 3 years, 2019, 2020 and 2021. Transgressive segregation was observed in all the three years.

To evaluate the stability of the DTH trait across generations, the F3 progeny of early- and late-flowering F2 individuals (EB and LB) were grown during Kharif 2020. Statistical analysis using Welch’s *t*-test revealed no significant differences in heading date between the F2 parents and their F3 progeny ***(Supplementary data S4)***, indicating that the trait is heritable and stable. This stability across generations highlights the potential of these segregants for breeding programs aimed at optimizing heading date and improving crop performance.

### QTL-seq Analysis Deciphered *qDTH3*, a High-Confidence Heading date QTL

The alignment of bulk and parental DNA reads to the reference genome enabled the calculation of the Δ SNP-index at each SNP position using the QTL-seq pipeline. Statistical confidence intervals were established to strengthen the identification of these genomic regions across the different seasons. A stretch of positive Δ SNP-index (peak) values crossing the 95% confidence interval (CI) was observed in Chr3 from 26.70-32.16 Mb ***(Supplementary data S5, Figure S2)*** in Kharif 2019, while the same chromosome exhibited a > 95% confidence peak interval from 26.96-32.29 Mb ***(Supplementary data S6, Figure S3)*** and 20.84-32.29 Mb ***(Supplementary data S7, Figure S4)*** in the Kharif 2020 and Kharif 2021 populations, respectively. Additionally, Chrs 6 (∼5 Mb and ∼25 Mb) and 7 (∼4 Mb) displayed stretches of negative Δ SNP-index (troughs) in the Kharif 2019 ***(Supplementary data S5, Figure S2)*** population, while in Kharif 2020 ***(Supplementary data S6, Figure S3)***, Chr6 (∼2-11 Mb) and 9 (10.5-16 Mb) displayed a trough and peak, respectively. In the Kharif 2021 ***(Supplementary data S7, Figure S4)*** population, Chr6 (∼2-11.5 Mb) and 8 (∼7.5-8 Mb) exhibited a high-confidence region of negative Δ SNP- index values. It is important to note that peaks typically indicate an allelic bias towards the EB (higher SNP-indexEB), while troughs represent a bias towards the alleles of LB (higher SNP-indexLB). In other words, the peak regions correspond to the genomic sequences from EB and SM93, whereas the troughs originate from the LB and SM alleles. Hence, the Chr3 interval is representative of the sequence in SM93. A common interval with > 95% confidence was detected in Chr3 from 26.96-32.16 Mb (5.2 Mb), in the three populations tested. This interval, referred to as *qDTH3*, was selected for further investigation **(*Figure 4*, *Table 1, Supplementary data S8)***.

**Figure 4:**
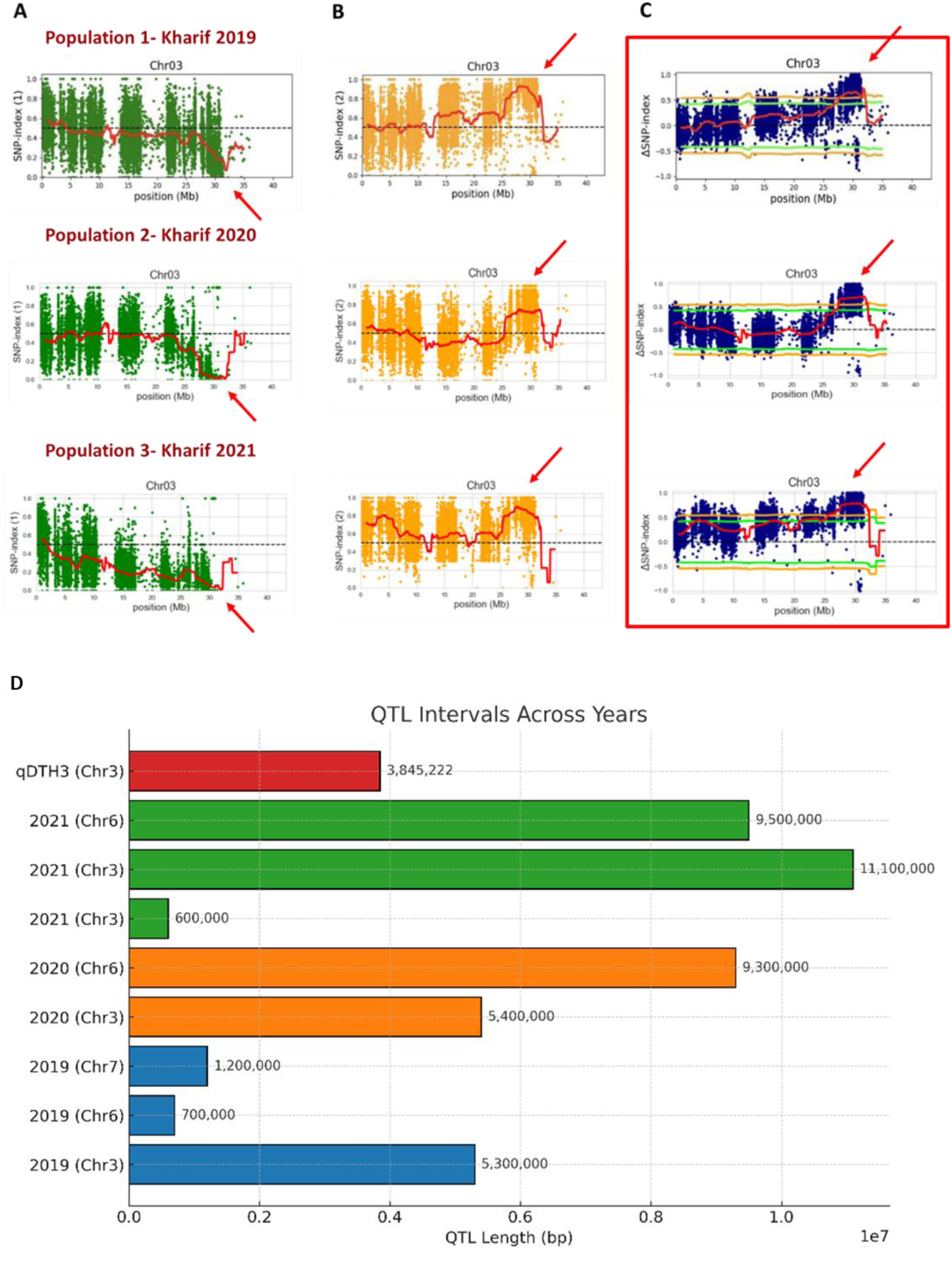
qDTH3 was Consistently Mapped in the Three Populations. **(A)** Chr3 SNP-index plot based on LB across three populations. **(B)** Chr3 SNP-index plot based on EB across three populations. **(C)** Chr3 Δ SNP-index Plot across three populations. **(D)** Length of QTL intervals detected in the 3 QTL-seq experiments.

**Table 1:** QTLs for early heading mapped in three different years.

| F2 Population | QTL Interval |  |  |  |
| --- | --- | --- | --- | --- |
|  | Chr | Start (bp) | End (bp) | QTL Length (bp) |
| 2019 | Chr3 | 27000000 | 32300000 | 5,300,000 |
|  | Chr6 | 4400000 | 5100000 | 700,000 |
|  | Chr7 | 2500000 | 3700000 | 1,200,000 |
| 2020 | Chr3 | 26900000 | 32300000 | 5,400,000 |
|  | Chr6 | 2000000 | 11300000 | 9,300,000 |
| 2021 | Chr3 | 4000000 | 4600000 | 600,000 |
|  | Chr3 | 21200000 | 32300000 | 11,100,000 |
|  | Chr6 | 2300000 | 11800000 | 9,500,000 |
| *qDTH3*(common QTL region in Chr3) | Chr3 | 27473190 | 31318412 | 3,845,222 |

It is worth noting that the Nipponbare IRGSP 1.0 genome assembly was employed for the QTL analyses, as the reference genome of SM was not available when this study was initiated. Subsequently, a reference genome for SM was generated (Rao et al., unpublished) and QTL mapping for early flowering in SM93 using this genome assembly has consistently produced similar results across the three populations tested with Nipponbare as the reference. **(*Figures S5, S6, S7*)**. Interestingly, the SM genome showed additional QTL intervals in other chromosomes at ∼5 Mb and ∼20 Mb in Chr12 (Kharif 2019, 2020 populations, respectively), ∼14-19 Mb and ∼20-22 Mb in Chr9 (Kharif 2020), ∼9-10 Mb in Chr8 (Kharif 2021). Further mapping and annotation studies are needed to confirm whether these QTLs are valid and reproducible.

### SNP Insights within *qDTH3*

The QTL intervals identified across the different Kharif seasons revealed a substantial number of SNPs, highlighting the extensive genetic diversity associated with heading date. In Kharif 2019, the identified QTL interval included 2736 SNPs, while Kharif 2020 yielded 2478 SNPs ***(Supplementary data S9, S10)***. Remarkably, the QTL analysis in Kharif 2021 uncovered an even larger set, with a total of 5578 SNPs ***(Supplementary data S11)***. Most strikingly, a core set of 2207 SNPs was found to be conserved across all three populations within *qDTH3*, underscoring their potential role in the genetic regulation of heading date ***(Figure 5A, Supplementary data S12)***. This core SNP set was linked to 2767 annotations, with half of these variants (1387) residing in genic regions. These SNPs overlapped with 217 genes and 313 transcripts ***(Supplementary data S13)***. Delving deeper into the genic regions, we discovered 221 SNPs in UTRs, 820 in introns, and 317 in coding regions of 123 genes, which were attributed to 153 transcripts ***(Figure 5, Supplementary data 14)***.

**Figure 5:**
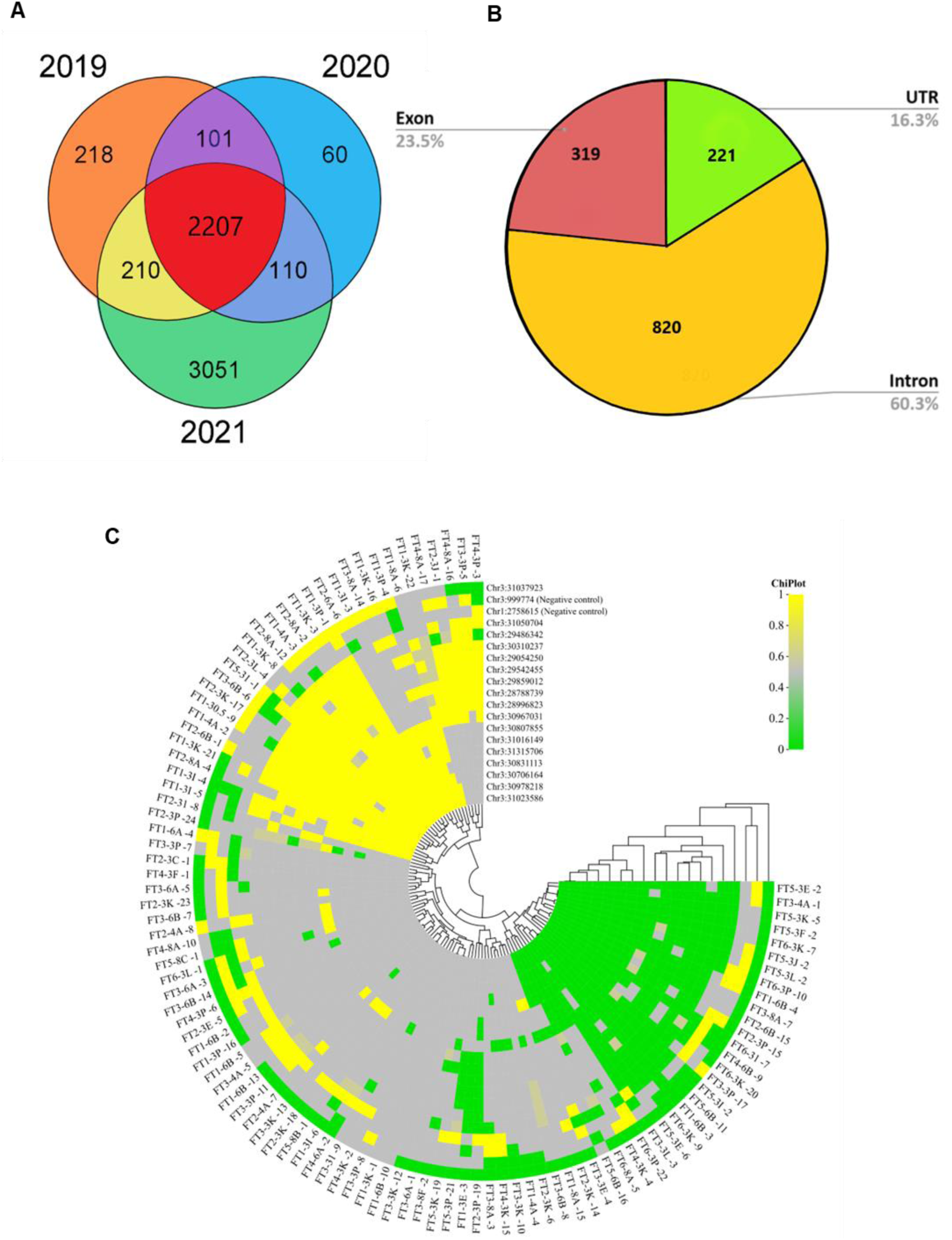
A**n**alysis **of SNPs associated with the early heading QTLs mapped in Chr3 (A)** Venn diagram showing the erlap of SNPs mapped in QTLs associated with early heading across three different years. **(B)** Classification of ic SNPs in qDTH3. **(C)** KASP analysis of SNPs in qDTH3. The phenotypes of the plants analyzed are indicated FT1-FT6, with FT1 being the earliest plants to flower, and FT6 the last.

### *qDTH3* Contains Multiple Genes Related to DTH

Analysis of *qDTH3* revealed that its SNPs overlapped with or were located near several QTLs and genes linked to growth and developmental processes, suggesting their potential role in regulating heading date. Specifically, these genes fell into four major categories: Flowering-related genes – This category includes transcription factors such as *OsMADS14*, *OsMADS34*, *OsSCL14*, and *OsCOL10*, which play key roles in floral transition. Additionally, other flowering-associated genes- *GF14f*, *OsWDR5a*, *OsWD40- 92*, and *TIP3 (TDR Interacting Protein 3)* were identified, each contributing to the transition to the reproductive stage, flower development, or both.

Phytohormone-related genes – These include the cytokinin receptor *HK4 (Histidine Kinase 4)* and a homolog of the Arabidopsis auxin-responsive gene *OS-QR (Quinone Reductase)*, both of which regulate hormonal signaling pathways affecting heading date.
Plant photoreceptors – The identification of *PhyA* and *PhyC*, two essential photoreceptors, suggests their involvement in light-dependent flowering regulation.
Genes involved in nutrient uptake and transport – Several genes within *qDTH3* were associated with nutrient signaling, including *OsAMT4 (Ammonium Transporter 4)*, *OsOPT7 (Iron-deficiency-regulated OligoPeptide Transporter 7)*, and *OsTPKA (Two-pore K+ channel family protein)*, potentially influencing DTH through metabolic regulation. The details of SNPs located in and around the aforementioned genes are provided in ***Table 2*** and ***Supplementary data S13***. Our investigation revealed that *qDTH3* also overlaps with rice QTLs reported for drought tolerance. We observed that the genes *OsITPK3 (Inositol 1,3,4-trisphosphate 5/6-kinase)* and *OsTPKA (Two-pore K^+^ channels)* located within *qDTH3* and harboring variants in SM93 are known to modulate osmotic stress tolerance in rice. In future, it would be interesting to delve deeper by conducting biochemical assays on SM93 to assess potential alterations in osmotic adjustment.

**Table 2:** SNP-Trait Association of SNPs within qDTH3 and their Annotations.

| S. No. | SNP | Nucleotide in Cultivar |  |  | Distance from previous SNP (kb) | Association Mapping |  | Gene ID | Gene Harboring the SNP |  |  |
| --- | --- | --- | --- | --- | --- | --- | --- | --- | --- | --- | --- |
|  | Chr:Position | Nip^ | SM | SM93 |  | p-value | R<sup>2</sup> value |  | Gene | Importance of the Gene | Variant Annotation |
| 1. | Chr3:999774 | T | A | T | - | 0.375 | 0.007 | Negative Controls for KASP genotyping |  |  |  |
| 2. | Chr1:2758615 | T | C | T | - | 0.146 | 0.019 |  |  |  |  |
| 3. | Chr3:28996823 | C | G | C | - | 6.43E-07 | 0.212 | Os03g0716200 | *TIP3 TDR Interacting Protein 3* | Anther development (Yang et al., 2019) | Upstream gene variant |
| 4. | Chr3:29054250 | G | A | G | 57.4 | 1.11E-06 | 0.204 | Os03g0717700 | *HK4 Histidine Kinase 4* | His-kinase domain (HK) of cytokinin receptor (Choi et al., 2012) | Missense variant in exon 4 causing a change M to I within CHASE domain for cytokinin binding, SIFT deleterious (0.03) |
| 5. | Chr3:29486342 | G | T | G | 432 | 4.38E-07 | 0.218 | Os03g0725300 | *OsWDR5a or DBR1 Lariat debranching enzyme* | Part of COMPASS-like complex, which modulates flowering (Jiang et al., 2018) | Missense variant in exon 1 of Os03t0725400-01 and Os03t0725400-03 and exon 2 of Os03t0725400-02, causing a G to C change. SIFT deleterious (0) |
| 6. | Chr3:29542455 | G | G | A | 56 | 1.72E-05 | 0.163 | Os03g0726200 | *OsITPK3 Inositol pyrophosphate kinase* | Drought and salt stresses (Du et al., 2011) | Missense variant in exon 1 causing S to N amino acid change. SIFT deleterious (0) |
| 7. | Chr3:29859012 | T | T | A | 316 | 4.92E-07 | 0.214 | Os03g0730400 | *OsSCP19 Similar to Serine carboxypeptidase* | - | Start lost M to K in exon 1 of Os03t0730400-01 and Os03t0730400-02. SIFT deleterious (0) |
| 8. | Chr3: 30310237 | G | G | T | 451 | 5.99E-07 | 0.213 | - | - | - | Intergenic variant |
| 9. | Chr3:30706164 | A | G | A | 847 | 5.55E-11 | 0.337 | Os03g0746800 | *OsWD40-92 tRNA (guanine-N(7))-methyltransferase non-catalytic subunit* | OsWD40 genes are involved in early developmental stages in seedlings, flower development (Ouyang et al., 2012) | Upstream gene variant |
| 10. | Chr3:30807855 | G | G | A | 101 | 1.08E-09 | 0.299 | Os03g0748400 | *OS-QR Quinone reductase* | Homologous to FQR1 in Arabidopsis (auxin responsive gene) that protects plant cells from oxidative stress (Laskowski et al., 2002) | Missense variant in exon 2 causing an A to V change, SIFT deleterious (0.02) |
| 11. | Chr3:30831113 | T | C | T | 23 | 2.96E-11 | 0.344 | Os03g0748900 | *OsAMT4 Ammonium Transporter* | OsAMTs are involved in ammonium uptake (Li et al., 2009) | Missense variant in exon 2 causing a V to A change in the ammonium transporter domain |
|  |  |  |  |  |  |  |  | Os03g0749051 | Hypothetical protein | - | 5 prime UTR variant |
| 12. | Chr3:30967031 | C | C | T | 135 | 3.84E-07 | 0.218 | Os03g0751100 | *OsOPT7 Iron-deficiency-regulated OligoPeptide Transporter 7* | Mobilization of organic nitrogenous compounds (Bashir et al., 2015) | Upstream gene variant |
| 13. | Chr3:30978218 | T | A | G | 11 | 6.68E-12 | 0.362 | Os03g0751400 | - | - | Upstream gene variant |
| 14. | Chr3:31016149 | G | G | A | 37 | 4.81E-10 | 0.309 | Os03g0752300 | *OsTPKA Two-pore K+ channel family protein* | Transgenic plants with the allele from a salt tolerant variety showed growth advantage in control and salt stress conditions (Haque et al., 2023). | Missense variant in exon 2 causing a R to W change, SIFT deleterious (0.02) |
| 15. | Chr3:31023586 | G | G | A | 7 | 6.68E-12 | 0.362 | Os03g0752700 | *OsDjC33 DnaJ domain protein C33* | - | Upstream gene variant |
| 16. | Chr3:31037923 | G | G | A | 14 | 2.09E-04 | 0.123 | Os03g0752800 | *OsMADS14* | Determination of floral meristem (Komiya et al., 2008). | Intron variant (2transcripts) |
| 17. | Chr3:31050704 | G | G | A | 12 | 2.77E-04 | 0.118 | Os03g0753100 | *OsMADS34/PAP2* | Specify the identity of the IM downstream of the florigen signal (Kobayashi et al., 2012). | Splice region variant, synonymous variant (exon 3) |
| 18. | Chr3:31315706 | T | T | C | 265 | 7.32E-11 | 0.333 | Os03g0757900 | *OsUGD3 UDP-glucose 6-dehydrogenase 3* | - | Upstream gene variant |

^Nipponbare

### SNP-trait Association Analysis Demonstrated a Significant Relationship between SNP Markers and DTH

Kompetitive Allele-Specific PCR (KASP^TM^) is a high-throughput genotyping method used for efficiently screening SNPs. In this study, we utilized KASP^TM^ to investigate 2,385 common SNPs identified across three mapping populations within *qDTH3*. By prioritizing high-confidence SNPs, particularly those with high Δ SNP values, and/or those associated with developmental and flowering genes, we refined our dataset to 47 SNPs designated for KASP assays ***(Supplementary data S15)***. 2 other SNPs not associated with the phenotype were used as negative controls; one from Chr1 and another from Chr3 (outside *qDTH3*). This selection included test SNPs from *qDTH3* as well as control SNPs from other regions, allowing us to explore genotype-trait associations and further our understanding of the genetic basis for variations in heading date. Firstly, 47 SNPs were screened in SM and SM93, after which 38 of them were picked, which showed a clear nucleotide difference between the two ***(Supplementary data S16)***. Further, we tested our EB and LB plants from the Kharif 2019 population for the 38 SNPs and screened for those SNPs that were consistent with the EB ***(Supplementary data S17)***. Finally, all the F2 individuals were genotyped for 18 SNPs (2 controls and 16 test) using KASP^TM^ assays ***(Supplementary data S18)***.

The initial efforts to screen 49 KASP^TM^ SNPs helped in arriving at 16 SNPs, thus narrowing the investigation to a 2.53 Mb (Chr3:28.78-31.31 Mb) interval within *qDTH3* (= 5.2 Mb, Chr3:26.96-32.16 MB). The association studies based on KASP^TM^ genotyping revealed a strong association between the various KASP^TM^ test SNPs and the phenotype, with *p*-values ranging from 6.68E-12 to 2.77E-04 ***(Table 2)***. In contrast, the negative controls, KASP1 and KASP2 displayed higher *p*-values of 0.375 and 0.146, respectively, clearly indicating a lack of association with the phenotype. A regression analysis of the associated SNPs provided R^2^-values ranging from ∼12% to 36% in the 2.53 Mb region. Collectively, the SNPs associated with heading date within *qDTH3* explain a quarter (mean = ∼25%) of the variation in the days to heading phenotype.

### Analysis Reveals Skewed Genome-wide Downregulation with Skewed Upregulation in *H3*

Panicle Initiation (PI) stage of the plants in Kharif 2020 was identified after multiple rounds ag leaf sheath sampling and dissection. PI of SM93 and SM occurred in a staggered fashion at 1 and 100 ±1 days after sowing, respectively. Isolation of reproductive shoot apical meristem es ***(Figure S8)*** at the respective PI stages and differential gene expression (DGE) analysis een the triplicates of SM93 and SM revealed widespread differential expression across the me. The read statistics, raw and normalized counts are described in ***(Supplementary data S19, S21)***. Out of 37,863 genes annotated in Nipponbare, 31,782 genes were initially identified as rentially expressed in our datasets, with 6911 genes meeting the filtering criteria for ficance (*p*-value ≤ 0.05, and log2 fold-change (log2FC) ≥ 1 for upregulation or log2FC ≤ -1 ownregulation) ***(Supplementary data S21, S22)***. Post filtering, there were 2865 upregulated s and 4045 downregulated genes in our dataset ***(Table 3*, *Figure 6A, Supplementary data S23)***. Here, ‘upregulation’ refers to higher expression in SM93 compared to SM. Similarly, ord ‘downregulation’ is used in the context of lower expression in SM93 compared to that in These results indicate a predominant trend of downregulation in SM93 in the genome-wide ext. Contrastingly, Chr3 had the highest proportion of upregulated genes (8.55%), calculated e number of upregulated genes divided by the total number of genes on the chromosome ressed in percentage) ***(Supplementary data S23)***. A closer look at Chr3, within the region in *H3* narrowed down by KASP^TM^ analysis showed that the upregulated genes slightly umbered the downregulated ones, with 45 genes showing increased expression and 43 ing reduced expression in SM93 ***(Figure 6B, Supplementary data S24)***. In essence, the DGE ysis highlights a predominant trend of genome-wide downregulation in SM93, with a localized ption on Chr3, where slight upregulation of genes in the *qDTH3* region may indicate a pensatory or regulatory role of *qDTH3* in controlling the heading date of SM93.

**Figure 6:**
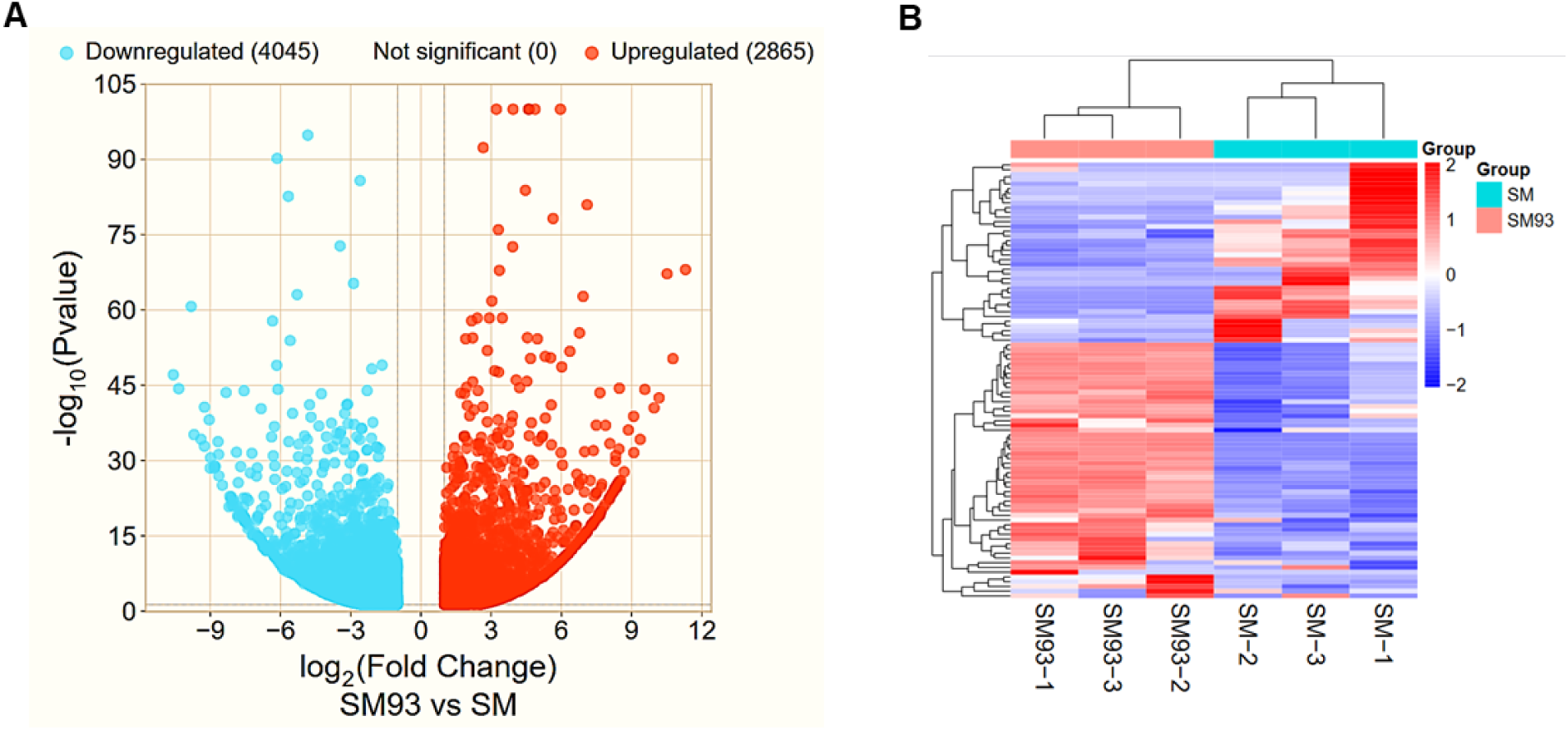
D**i**fferential **gene expression analysis of SM93 vs SM (A)** 2865 genes and 4045 genes were found to be Upregulated and downregulated globally in SM93 respectively. **(B)** Normalised counts of genes within refined H3 are slightly skewed towards upregulation.

**Table 3:** Differential gene expression analysis of SM93 vs SM is indicative of an overall downregulation of genes 93.

| Gene Category | Raw | Filtered |
| --- | --- | --- |
| Upregulated genes | 14745 | 2865 |
| Downregulated genes | 17037 | 4045 |
| Differentially expressed genes | 31782 | 6911 |
| Total genes studied | 37863 |  |

### Several *MADS-box* Genes were Upregulated in SM93

Gene Ontology (GO) enrichment analysis of filtered DEGs performed with RiceFREND, h integrates microarray data from RiceXPro, revealed a significant association of the rentially expressed genes (DEGs) with GO:0003700 (transcription factor activity). ifically, 134 genes were mapped to this GO term ***(Figure 7A, Supplementary data S25)***. To plement this, we conducted another GO enrichment analysis using gProfiler, which revealed ional enrichment of other biological processes, including GO:0042221 (response to ical). Specifically, 367 genes were linked to chemical response processes, and 298 genes associated with DNA-binding transcription factor activity ***(Figure 7B, Supplementary data S26)*** . Within the enriched set, the *qDTH3* candidate region notably showed upregulation of *zinc r homeodomain class homeobox transcription factor 11 (OsZHD11, Os03g0718500)*, *heat s transcription factor A2A (HSFA2A, Os03g0745000)*, and *panicle phytomer 2/MADS-box 34 (OsMADS34/PAP2*, *Os03g0753100)*, and downregulation of *Homeobox protein knotted- e 7 (OsH3, Os03g0727200)*, and, *Homeobox protein knotted-1-like 6 (OSH1/Oskn1*, *g0727000)*. Additionally, a gene set enrichment analysis using the PAGE tool on AgriGO, h took into consideration the fold changes in gene expression, indicated an upregulation of GO term for reproductive process (GO:0048608) ***(Figure 7C, Supplementary data S27)***. A led analysis of the genes categorized under this GO term threw light on the upregulation of iple transcription factors, particularly, the MADS-box genes. We conducted a search for genes ciated with this GO term and discovered that 19 of them exhibited upregulation, whereas 7 s displayed downregulation, and hence the GO term showed an overall upregulation ***(Figure Table 4)***. Interestingly among these DEGs, 11 MADS-box transcription factor genes were d to be upregulated, while 3 of them were downregulated. Particularly, E-class genes, which omologs of *Arabidopsis SEPALLATA (SEP)* were highly upregulated. Namely, *OsMADS1* old upregulation in SM93), *OsMADS5* (∼80-fold upregulation in SM93), *OsMADS6* (23.5- upregulation in SM93), *OsMADS7* (4.75-fold upregulation in SM93), *OsMADS17* (>6-fold gulation in SM93), *OsMADS24* (>5.5-fold upregulation in SM93), *OsMADS34* (∼100-fold gulation in SM93). The GO terms GO:0006082 (organic acid metabolic process), 0006633 (Fatty acid biosynthetic process) and GO:0006520 (Cellular amino acid biosynthetic ess) were enriched and showed an overall upregulation in SM93, which highlights enhanced bolic efficiency in SM93. The results from the GO enrichment and gene set enrichment yses suggest that the early heading phenotype observed in the SM93 mutant is likely enced by the interplay of transcriptional regulation and chemical signaling pathways, ding potential roles for phytohormones and their downstream regulatory networks.

**Figure 7:**
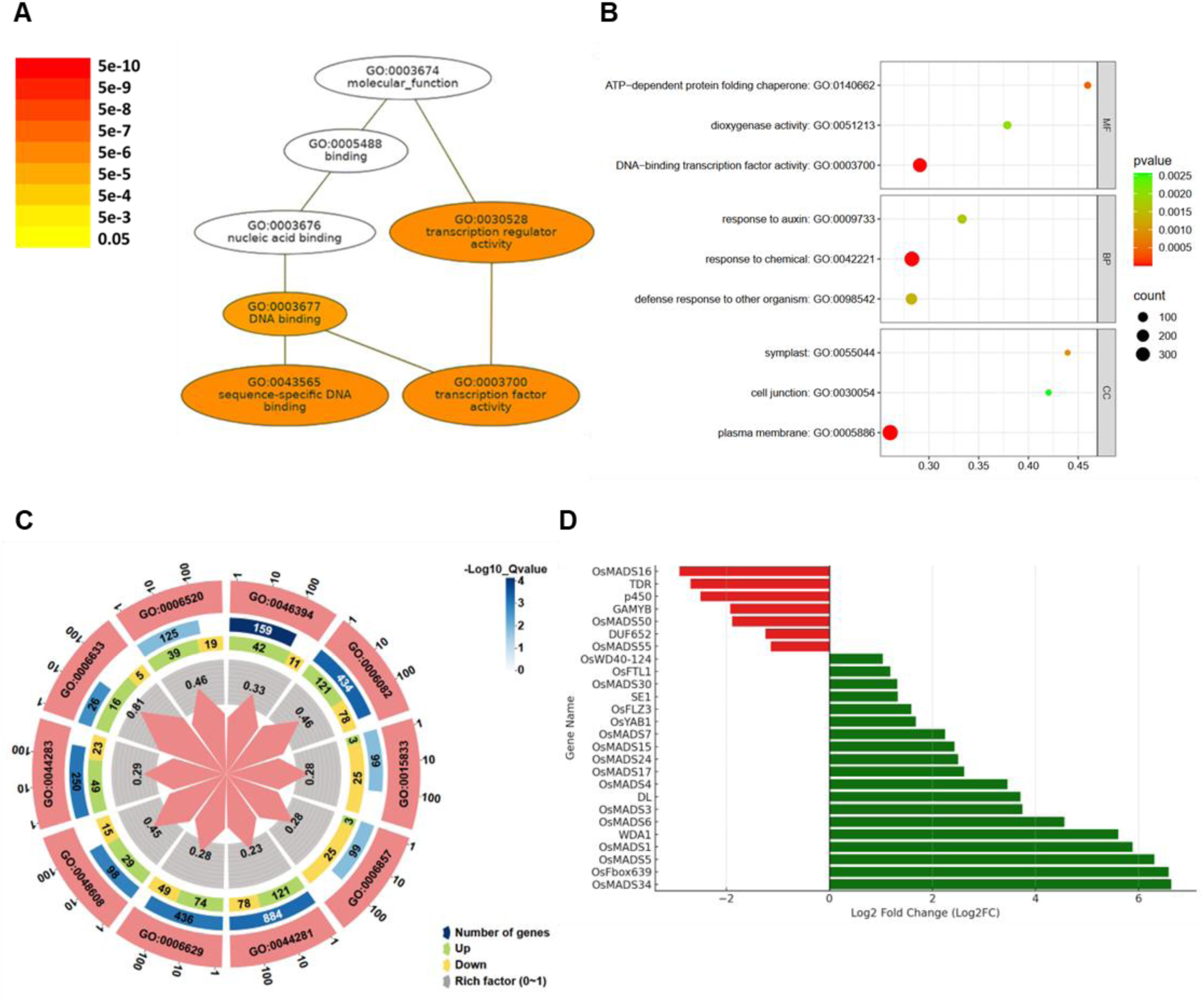
D**E**Gs **found in the reproductive SAM analysis between SM93 and SM. (A)** Bubble chart showing the enriched GO terms based on gProfiler analysis. Enriched terms include DNA-binding transcription factor ty, response to chemical, and response to auxin, with bubble sizes representing the number of genes associated each term and colors indicating the enrichment significance. **(B**) PAGE analysis using AgriGO indicates that 006082 (organic acid metabolic process), GO:0048608 (Reproductive process), GO:0006633 (Fatty acid nthetic process), GO:0006520 (Cellular amino acid biosynthetic process) are enriched and show an overall ulation in SM93. **(D)** Fold changes in genes involved in reproductive structure development.

**Table 4:** DEGs Attributed to Reproductive Structure Development.

| S. No. | Gene ID | Gene name | Gene Name | log2FC |
| --- | --- | --- | --- | --- |
| 1. | Os03g0753100 | *OsMADS34, PAP2* | *MADS-box transcription factor 34, Panicle Phytomer 2* | 6.64 |
| 2. | Os12g0129700 | *OsFbox639* | *Cyclin-like F-box domain containing protein* | 6.59 |
| 3. | Os06g0162800 | *OsMADS5* | *MADS-box transcription factor 5* | 6.31 |
| 4. | Os03g0215400 | *OsMADS1, LHS1* | *MADS-box transcription factor 1, Leafy Hull Sterile 1* | 5.89 |
| 5. | Os10g0471100 | *WDA1* | *Wax-Deficient Anther 1* | 5.61 |
| 6. | Os02g0682200 | *OsMADS6, MFO1, AFG1* | *MADS-box transcription factor 56, Mosaic Floral Organs 1* | 4.56 |
| 7. | Os01g0201700 | *OsMADS3, RAG* | *MADS-box transcription factor 3, Rice Agamous* | 3.75 |
| 8. | Os03g0215200 | *DL* | *Drooping leaf-1, DL protein* | 3.71 |
| 9. | Os05g0423400 | *OsMADS4* | *MADS-box transcription factor 4, Rice PISTILLATA ortholog* | 3.46 |
| 10. | Os04g0580700 | *OsMADS17* | *MADS-box transcription factor 17* | 2.62 |
| 11. | Os09g0507200 | *OsMADS24* | *MADS-box transcription factor 8* | 2.50 |
| 12. | Os07g0108900 | *OsMADS15, DEP* | *MADS-box transcription factor 15, Degenerative palea* | 2.43 |
| 13. | Os08g0531700 | *OsMADS7* | *MADS-box transcription factor 7, AGAMOUS-like 6* | 2.25 |
| 14. | Os07g0160100 | *OsYAB1* | *YABBY1* | 1.68 |
| 15. | Os01g0719000 | *OsFLZ3* | *FCS-LIKE ZINC FINGER PROTEIN 3* | 1.59 |
| 16. | Os06g0275000 | *SE1, Hd1* | *Heading date 1, Photosensitivity1* | 1.33 |
| 17. | Os06g0667200 | *OsMADS30* | *MADS-box transcription factor 30* | 1.32 |
| 18. | Os01g0218500 | *OsFTL1* | *FLOWERING LOCUS T (FT)-Like homolog* | 1.18 |
| 19. | Os06g0131100 | *OsWD40-124* | *Similar to guanine nucleotide-binding protein beta subunit-like protein 1* | 1.04 |
| 20. | Os06g0217300 | *OsMADS55* | *MADS-box transcription factor 55* | -1.14 |
| 21. | Os02g0468200 | *DUF652 family protein* | *DUF652-like unknown protein* | -1.24 |
| 22. | Os03g0122600 | *OsMADS50* | *MADS-box transcription factor 50* | -1.89 |
| 23. | Os01g0812000 | *GAMYB* | *Transcriptional activator of gibberellin-dependent alpha-amylase expression* | -1.93 |
| 24. | Os03g0134500 | *p450* | *Cytochrome p450* | -2.5 |
| 25. | Os02g0120500 | *TDR* | *Tapetum degeneration retardation, basic helix loop helix 5* | -2.70 |
| 26. | Os06g0712700 | *OsMADS16, SPW1* | *MADS-box transcription factor 16, Superwoman-1* | -2.91 |

### Integration of QTL Mapping and RNA-Seq Data Reveals Candidate Genes y

A key *MADS-box* transcription factor, *Panicle Phytomer 2/MADS-box gene 34 (Os03g0753100, PAP2/OsMADS34)*, located within the refined qDTH3 interval, exhibited the highest expression in SM93, approximately 100 times higher compared to SM. A synonymous SNP (exonic, splice region variant) in this gene was strongly associated with the trait in our KASPTM assay (***Table 2***). This gene contains 4 intronic SNPs. The cause of OsMADS34 upregulation remains unclear at this time, and the splice region SNP did not seem to cause any apparent splicing variation as per our RNA-seq data. The gene, HK4 (Os03g0717700) was significantly downregulated, and a SNP-trait association were established for a missense variant in exon 4 of HK4, which likely results in a methionine (M) to isoleucine (I) substitution within the CHASE domain, responsible for cytokinin binding. SIFT analysis predicted this change to be deleterious, potentially disrupting protein function. However, HK4 harbors 12 SNPs in total, with 3 located upstream of the gene at -420, -335, and -256 bp, which may be part of cis-elements, contributing to the candidate SNP markers for the heading date QTL identified in this study. Similarly, OsSCP19 (Os03g0730400), a serine carboxypeptidase, contains a missense mutation that likely causes the loss of the start codon. This SNP was strongly associated with the early heading phenotype in SM93, and the gene was significantly downregulated in our data (***Table 2***). Among the 7 SNPs identified in OsSCP19, 2 were located -67 and -8 bp upstream of the gene, which may also be part of cis-elements. Serine carboxypeptidases are known to respond to phytohormones, and GS5, a serine carboxypeptidase, in particular, promotes mitosis in young panicle tissues, thereby controlling grain size (Washio & awa, 1994; Xu et al., 2015), it remains unclear whether OsSCP19 plays a role in the regulation of heading date in rice, or if its association with early heading is due to linkage with other key genes within the QTL region, and hence our findings warrant further investigation.

## DISCUSSION

### Transgressive DTH Phenotypes Broaden the Trait Distribution

Transgressive segregation is a common occurrence in the context of DTH phenotypes (Hori et al., 2015; Thomson et al., 2006; Yan et al., 2011). Epistasis and gene interaction often lead to transgressive phenotypes, wherein novel combinations of complementary alleles are created during hybridization (Z. Li et al., 1995; Thomson et al., 2006). The observation that the F1 hybrids of SM x SM93 flowered even earlier than SM93 by ∼10 days is interesting and could be due to a complementation of recessive alleles or heterozygosity at heading date controlling loci including *qDTH3* or some other reasons. The difference between the DTH of SM and SM93 was about 7-8 days. However, the DTH variation in the F2 population of their cross ranged from 36 days in Kharif 2019, to 42 days in Kharif 2020 and 2021. The variation in DTH between the different years could be due to environmental effects. The genetic basis of the extreme DTH phenotypes however, is evident from the heritability of the extreme traits in the F3 generation. Z. Li et al. (1995) also observed a marked transgressive segregation in the F2 progeny of a cross between the parental lines, ’Lemont’ and ’Teqing’ rice varieties, which have very similar heading dates (Z. Li et al., 1995). They have suggested that both the parents contained alleles for early heading at different QTLs, which united in the early transgressive segregants (Z. Li et al., 1995). On similar lines, the Chr6 QTL (trough) obtained through our QTL-seq analysis with sequences corresponding to SM, may be partly responsible for the transgressive phenotype in our F2 population, wherein the underlying mechanism for transgressive segregation could be that of gene interaction between the *qDTH3* alleles (from SM93) and the alleles in the Chr6 QTL (similar to SM). Such progenies have broadened the scope of our future studies and provided candidates for more extreme phenotypes than SM93.

### *qDTH3* and Strong SNP-Trait Associations

The QTL-seq strategy employed to map the early flowering QTL in SM93 from three F2 mapping populations (SM x SM93) led us to a 5.2 Mb region on Chr3, which we refer to as *qDTH3*. The QTARO (http://qtaro.abr.affrc.go.jp/) database was looked up to compare the results of this study with known QTLs for DTH in rice. The QTLs that overlapped partly or completely with *qDTH3* were attributed to heading date, flowering time, drought tolerance, panicle traits and biotic stress tolerance in various studies. The QTL was refined to 2.53 Mb using gene annotations, Δ SNP-index values, SIFT scores, and KASP genotyping. Multiple candidate genes were identified, including those related to nutrient uptake, phytohormones, and flowering. Sixteen candidate SNPs were linked to the phenotype, with two affecting functional domains- *HK4* (CHASE domain) and *OsAMT4* (ammonium transporter domain). SNPs were also found in *OsMADS14* and *OsMADS34*, key flowering regulators. *qDTH3* may comprise multiple sub-QTLs influencing vegetative growth, phytohormone response, and reproductive transition.

Reciprocal backcross inbreeding of the *japonica* rice cultivars Nipponbare and Koshihikari with a DTH variation of 7 days gave rise to transgressive DTH segregants in both the backcross inbred lines (BILs) (Matsubara et al., 2008). Two QTLs attributed to transgressive DTH segregants named, *Hd16* (1,140 kb) and *Hd17* (328 kb) were discovered in the same study in Chrs 3 and 6, respectively, based on SSR and SNP markers through conventional QTL mapping. These regions may overlap with the QTLs that have been detected in our study, but are yet to be fine-mapped for the candidate gene(s) and causal SNP(s) (Matsubara et al., 2008). Uga et al., 2007 described accumulation of additive effects of QTLs, leading to extremely late heading of Nona Bokra, an *indica* rice variety, under natural field (Tsukuba, Ibaraki, Japan) and long-day conditions (14.5 hours). One of the QTLs named as QTAROqtl-745 in Chr3 lies close to *qDTH3* (Uga et al., 2007). Under short day (10 hours) conditions, Nona Bokra showed extremely early heading.

Genetic background influences trait variation, making it difficult to integrate results across studies, especially for complex traits like heading date, which involve epistatic interactions.

Interactions between loci are stronger in *indica* than *japonica* rice, with *indica* having more haplotypes for heading date genes (Liu et al., 2021). It was intriguing that a Genome-Wide Association Study (GWAS) on 278 *indica* and 147 *japonica* accessions uncovered distinct QTLs associated with heading date in *japonica* and *indica* rice, under both Short-Day (SD) and Long-Day (LD) conditions. *Japonica* rice varieties grown in temperate regions have developed weaker photoperiod sensitivity compared to *indica* rice, which is more sensitive to day length changes typical of tropical regions. Hence, the subpopulation-associated distinctness can be attributed to the weak photoperiod sensitivity in *japonica* rice under SD conditions (Han et al., 2016). There is an abundance of haplotypes within heading date genes in *indica* rice, ranging from non-functional to weak and strong responses to photoperiod.

### Upregulation of the *MADS-box* Gene Network

DGE of various *MADS-box* transcription factors between SM and SM93 was observed at the panicle initiation stage. Interestingly, the refined *qDTH3* interval contains *OsMADS34*, which exhibits high upregulation. Furthermore, a SNP in *OsMADS34* was significantly associated with early heading in our KASP assay (*p*-value = 2.77E-04). *OsMADS34/PAP2* plays a role in specifying Inflorescence Meristem (IM) development downstream of the rice florigens, and in promoting the transition from branch meristems to spikelet meristems (Gao et al., 2010; Kobayashi et al., 2012). *OsMADS5*, *OsMADS1* and *OsMADS34* interact with each other, and with other MADS-box floral homeotic members, and together regulate floral meristem determinacy. They specify the identities of spikelet organs by positively regulating the other MADS-box floral homeotic genes (Wu et al., 2018). *OsMADS6* is involved in floral meristem determination, along with *OsMADS17,* (Dong et al., 2023; H. Li et al., 2010). *OsMADS7* is involved in determining flowering time, and homeotic functions, along with *OsMADS24* (Cui et al., 2010).

### Candidate Cytokinin and GA-Related Genes in Modulating Early Heading

Cytokinin is a plant hormone that regulates meristem function and promotes cell division, impacting plant growth and development. Changes in cytokinin levels can indirectly affect heading date (Cho et al., 2022). In our study, we identified a cytokinin (CK) receptor gene, *Histidine Kinase 4 (HK4)*, located within the refined *qDTH3* region, which was significantly associated with the early heading phenotype in our KASP assay. The expression of *HK4* was significantly downregulated in SM93. A missense mutation (M→I) in the CHASE domain (extracellular cytokinin-binding domain) of *HK4* was predicted to be deleterious, suggesting a disruption in receptor function. Mutant plants lacking cytokinin receptors exhibited delayed bolting in *Arabidopsis* (Nishimura et al., 2004; Riefler et al., 2006). However, another study suggests that externally applied cytokinin can postpone heading date and extend the vegetative phase by suppressing florigen (Cho et al., 2022). This highlights the complex role of cytokinin in heading date regulation, where both receptor signaling and external cytokinin levels may have opposing effects, depending on the context, such as the plant’s internal hormonal balance or timing of exposure.

In addition to *HK4*, we observed the downregulation of KNOX (KNOTTED-like homeobox) transcription factors (TFs), such as *OSH1* and *OSH3*, in SM93. KNOX TFs are key regulators of cytokinin biosynthesis in meristems (Sakamoto et al., 2006), and their downregulation could further contribute to altered cytokinin homeostasis in this variety. Studies have shown that KNOX TFs not only promote cytokinin biosynthesis but also regulate gibberellin (GA) catabolism, suggesting a tight interdependence between GA and cytokinin signaling pathways. In our study, we found that both GA biosynthesis genes, such as *GNP1 (GA20ox1)*, and GA catabolism genes (e.g., *GA2ox1*, *GA2ox3*, *GA2ox4*) were downregulated. However, the overall levels of GA and cytokinin in SM93 remain uncertain, and further estimation of hormone content is necessary to determine whether the levels are indeed reduced or increased, as the gene expression data alone does not confirm the direction of these hormonal changes. The interplay between GA and cytokinin is complex and context-dependent. While cytokinin generally promotes vegetative growth and meristem activity, GA is known to promote flowering and heading in rice (Yamaguchi & Kamiya, 2000). Are changes in GA biosynthesis and/or catabolism, coupled with alterations in cytokinin biosynthesis and/or signaling, driving a disruption in hormonal homeostasis in SM93, thereby influencing the timing of flowering by affecting the shift from vegetative to reproductive growth? We hypothesize that the reduced expression of *HK4* in SM93, coupled with the downregulation of KNOX TFs, disrupts the cytokinin-GA balance, leading to altered heading date. These changes, potentially exacerbated by the mutations in the *HK4* receptor and its altered binding activity, may underlie the observed early heading phenotype in SM93. Future studies involving hormone quantification and genetic manipulation of KNOX and cytokinin signaling genes will provide deeper insights into the hormonal regulation of heading date in rice.

## CONCLUSIONS

This study identifies a critical QTL associated with the early flowering, providing significant insights into its genetic basis. The candidate SNPs, validated through genotyping, exhibit a robust association with early flowering, confirming their potential as molecular markers. Transcriptomic analysis further reveals that key genes within the *MADS-box* gene network are markedly upregulated in SM93, suggesting a possible role for them in regulating heading date. In addition, a cytokinin-gibberellin (CK-GA) rebalancing mechanism may potentially underpin its early heading. The *qDTH3* region, which accounts for approximately 25% of the trait variance, exerts a profound genetic effect on the early flowering phenotype. To optimize the application of these findings, fine mapping of the *qDTH3* region is essential to minimize the risk of undesirable linked traits before implementing SNP-based plant screening and QTL introgression into breeding programs.

## ACKNOWLEDGEMENTS

We sincerely acknowledge the Department of Biotechnology (DBT), Ministry of Science and Technology, Government of India, for funding under the Category-I Biotechnology Research Fellowships Program provided to DR. This study was also supported by grants to MSM, HKP, and RVS from the Council of Scientific and Industrial Research (CSIR), Ministry of Science and Technology, Government of India (MLP0121 Phase-I and Phase- II, MLP0057) and the JC Bose Fellowship of RVS granted by the Science and Engineering Research Board (SERB), Department of Science and Technology, Government of India (SB/S2/JCB-12/2014). Additionally, we appreciate the skilled workers at CSIR-CCMB for their support in field activities and maintenance.

## AUTHOR CONTRIBUTION STATEMENT

HKP, RVS: Conceptualization, supervision, project administration, manuscript review and editing, funding acquisition. MSM, RMS: provided seed material, suggestions for mapping studies; DR, GCG: Methodology. DR: Cross-pollination, sample collection, investigation (phenotyping, QTL-seq, RNA-seq analysis), formal analysis, and writing the original draft. NST: Data entry, KASP genotyping, and assistance with investigation (phenotyping, sample collection, QTL-seq, RNA-seq analysis). GCG, NG, MJ: Cross- pollination. MJ: Field maintenance. All authors read and approved the final manuscript.

## DECLARATIONS

### Ethics approval

Not applicable

### Consent to participate

Not applicable

### Consent to publish

Not applicable

## REFERENCE

Bashir, K., Ishimaru, Y., Itai, R. N., Senoura, T., Takahashi, M., An, G., Oikawa, T., Ueda, M., Sato, A., Uozumi, N., Nakanishi, H., & Nishizawa, N. K. (2015). Iron deficiency regulated OsOPT7 is essential for iron homeostasis in rice. Plant Molecular Biology, 88(1–2), 165–176. 10.1007/s11103-015-0315-0

Cho, L. H., Yoon, J., Tun, W., Baek, G., Peng, X., Hong, W. J., Mori, I. C., Hojo, Y., Matsuura, T., Kim, S. R., Kim, S. T., Kwon, S. W., Jung, K. H., Jeon, J. S., & An, G. (2022). Cytokinin increases vegetative growth period by suppressing florigen expression in rice and maize. Plant Journal, 110(6), 1619–1635. 10.1111/tpj.15760

Choi, J., Lee, J., Kim, K., Cho, M., Ryu, H., An, G., & Hwang, I. (2012). Functional identification of oshk6 as a homotypic cytokinin receptor in rice with preferential affinity for iP. Plant and Cell Physiology, 53(7), 1334–1343. 10.1093/pcp/pcs079

Cui, R., Han, J., Zhao, S., Su, K., Wu, F., Du, X., Xu, Q., Chong, K., Theißen, G., & Meng, Z. (2010). Functional conservation and diversification of class e floral homeotic genes in rice (Oryza sativa). Plant Journal, 61(5), 767–781. 10.1111/j.1365-313X.2009.04101.x

Dong, H., Qiu, Y., Wang, T., Song, S., Li, L., & Li, Y. (2023). Functional identification of OsMADS6 and OsMADS17 in floral organ development in rice. Plant Breeding. 10.1111/pbr.13103

Doyle, J., & Hortorium, B. (1991). DNA PROTOCOLS FOR PLANTS CT AB Total DNA Isolation.

Du, H., Liu, L., You, L., Yang, M., He, Y., Li, X., & Xiong, L. (2011). Characterization of an inositol 1,3,4-trisphosphate 5/6-kinase gene that is essential for drought and salt stress responses in rice. Plant Molecular Biology, 77(6), 547–563. 10.1007/s11103-011-9830-9

Gao, X., Liang, W., Yin, C., Ji, S., Wang, H., Su, X., Guo, C., Kong, H., Xue, H., & Zhang, D. (2010). The SEPALLATA-like gene OsMADS34 is required for rice inflorescence and spikelet development. Plant Physiology, 153(2), 728–740. 10.1104/pp.110.156711

Han, Z., Zhang, B., Zhao, H., Ayaad, M., & Xing, Y. (2016). Genome-wide association studies reveal that diverse heading date genes respond to short and long day lengths between Indica and Japonica rice. Frontiers in Plant Science, 7(AUG2016). 10.3389/fpls.2016.01270

Haque, U. S., Elias, S. M., Jahan, I., & Seraj, Z. I. (2023). Functional genomic analysis of K+ related salt-responsive transporters in tolerant and sensitive genotypes of rice. Frontiers in Plant Science, 13. 10.3389/fpls.2022.1089109

Hori, K., Nonoue, Y., Ono, N., Shibaya, T., Ebana, K., Matsubara, K., Ogiso-Tanaka, E., Tanabata, T., Sugimoto, K., Taguchi-Shiobara, F., Yonemaru, J. ichi, Mizobuchi, R., Uga, Y., Fukuda, A., Ueda, T., Yamamoto, S. ichi, Yamanouchi, U., Takai, T., Ikka, T., … Yano, M. (2015). Genetic architecture of variation in heading date among Asian rice accessions. BMC Plant Biology, 15(1). 10.1186/s12870-015-0501-x

Jiang, P., Wang, S., Jiang, H., Cheng, B., Wu, K., & Ding, Y. (2018). The COMPASS- like complex promotes flowering and panicle branching in rice. Plant Physiology, 176(4), 2761–2771. 10.1104/pp.17.01749

Kobayashi, K., Yasuno, N., Sato, Y., Yoda, M., Yamazaki, R., Kimizu, M., Yoshida, H., Nagamura, Y., & Kyozukaa, J. (2012). Inflorescence meristem identity in rice is specified by overlapping functions of three AP1/FUL-Like MADS box genes and PAP2, a SEPALLATA MADS Box gene. Plant Cell, 24(5), 1848–1859. 10.1105/tpc.112.097105

Komiya, R., Ikegami, A., Tamaki, S., Yokoi, S., & Shimamoto, K. (2008). Hd3a and RFT1 are essential for flowering in rice. Development, 135(4), 767–774. 10.1242/dev.008631

Komiya, R., Yokoi, S., & Shimamoto, K. (2009). A gene network for long-day flowering activates RFT1 encoding a mobile flowering signal in rice. Development, 136(20), 3443–3450. 10.1242/dev.040170

Laskowski, M. J., Dreher, K. A., Gehring, M. A., Abel, S., Gensler, A. L., & Sussex, I. M. (2002). FQR1, a novel primary auxin-response gene, encodes a flavin mononucleotide-binding quinone reductase. Plant Physiology, 128(2), 578–590. 10.1104/pp.010581

LI, B. zhen, MERRICK, M., LI, S. mei, LI, H. ying, ZHU, S. wen, SHI, W. ming, & SU, Y. hua. (2009). Molecular Basis and Regulation of Ammonium Transporter in Rice. Rice Science, 16(4), 314–322. 10.1016/S1672-6308(08)60096-7

Li, H., Liang, W., Jia, R., Yin, C., Zong, J., Kong, H., & Zhang, D. (2010). The AGL6- like gene OsMADS6 regulates floral organ and meristem identities in rice. Cell Research, 20(3), 299–313. 10.1038/cr.2009.143

Li, Z., Pinson, S. R. M., Stansel, J. W., Park, W. D., Hart, G. E., & Park, ([]) -W D. (1995). Identification of quantitative trait loci (QTLs) for heading date and plant height in cultivated rice (Oryza sativa L.). In Theor Appl Genet (Vol. 91). Springer-Verlag.

Liu, C., Tu, Y., Liao, S., Fu, X., Lian, X., He, Y., Xie, W., & Wang, G. (2021). Genome-wide association study of flowering time reveals complex genetic heterogeneity and epistatic interactions in rice. Gene, 770, 145353. 10.1016/j.gene.2020.145353

Mansfeld, B. N., & Grumet, R. (2018). QTLseqr: An R Package for Bulk Segregant Analysis with Next-Generation Sequencing. The Plant Genome, 11(2). 10.3835/plantgenome2018.01.0006

Matsubara, K., Kono, I., Hori, K., Nonoue, Y., Ono, N., Shomura, A., Mizubayashi, T., Yamamoto, S., Yamanouchi, U., Shirasawa, K., Nishio, T., & Yano, M. (2008). Novel QTLs for photoperiodic flowering revealed by using reciprocal backcross inbred lines from crosses between japonica rice cultivars. Theoretical and Applied Genetics, 117(6), 935–945. 10.1007/s00122-008-0833-0

Nishimura, C., Ohashi, Y., Sato, S., Kato, T., Tabata, S., & Ueguchi, C. (2004). Histidine kinase homologs that act as cytokinin receptors possess overlapping functions in the regulation of shoot and root growth in arabidopsis. Plant Cell, 16(6), 1365–1377. 10.1105/tpc.021477

Ouyang, Y., Huang, X., Lu, Z., & Yao, J. (2012). Genomic survey, expression profile and co-expression network analysis of OsWD40 family in rice. BMC Genomics, 13(1). 10.1186/1471-2164-13-100

Riefler, M., Novak, O., Strnad, M., & Schmülling, T. (2006). Arabidopsis cytokinin receptors mutants reveal functions in shoot growth, leaf senescence, seed size, germination, root development, and cytokinin metabolism. Plant Cell, 18(1), 40–54. 10.1105/tpc.105.037796

Sakamoto, T., Sakakibara, H., Kojima, M., Yamamoto, Y., Nagasaki, H., Inukai, Y., Sato, Y., & Matsuoka, M. (2006). Ectopic Expression of KNOTTED1-Like Homeobox Protein Induces Expression of Cytokinin Biosynthesis Genes in Rice 1[W]. Plant Physiology. 10.1104/pp.106.085811

Shojiro Tamaki, Shoichi Matsuo, Hann Ling Wong, Shuji Yokoi, & Ko Shimamoto. (2007). Hd3a Protein Is a Mobile Flowering Signal in Rice. Science, 316(5827), 1030–1036. 10.1126/science.1141752

Sugihara, Y., Young, L., Yaegashi, H., Natsume, S., Shea, D. J., Takagi, H., Booker, H., Innan, H., Terauchi, R., & Abe, A. (2022). High-performance pipeline for MutMap and QTL-seq. PeerJ. 10.7717/peerj.13170

Sun, K., Huang, M., Zong, W., Xiao, D., Lei, C., Luo, Y., Song, Y., Li, S., Hao, Y., Luo, W., Xu, B., Guo, X., Wei, G., Chen, L., Liu, Y. G., & Guo, J. (2022). Hd1, Ghd7, and DTH8 synergistically determine the rice heading date and yield-related agronomic traits. Journal of Genetics and Genomics, 49(5), 437–447. 10.1016/j.jgg.2022.02.018

Takagi, H., Abe, A., Yoshida, K., Kosugi, S., Natsume, S., Mitsuoka, C., Uemura, A., Utsushi, H., Tamiru, M., Takuno, S., Innan, H., Cano, L. M., Kamoun, S., & Terauchi, R. (2013). QTL-seq: Rapid mapping of quantitative trait loci in rice by whole genome resequencing of DNA from two bulked populations. Plant Journal, 74(1), 174–183. 10.1111/tpj.12105

Thomson, M. J., Edwards, J. D., Septiningsih, E. M., Harrington, S. E., & McCouch, S. R. (2006). Substitution mapping of dth1.1, a flowering-time quantitative trait locus (QTL) associated with transgressive variation in rice, reveals multiple sub-QTL. Genetics, 172(4), 2501–2514. 10.1534/genetics.105.050500

Vergara, B. S., Tanaka, A., Lilis, R., & Puranabhavung, S. (1966). Relationship between growth duration and grain yield of rice plants. Soil Science and Plant Nutrition, 12(1), 31–39. 10.1080/00380768.1966.10431180

Washio, K., & Ishikawa, K. (1994). Organ-Specific and Hormone-Dependent Expression of Genes for Serine Carboxypeptidases during Development and Following Germination of Rice Grains. In *Plant Physiol* (Vol. 105). www.plantphysiol.org

Won, P. L. P., Liu, H., Banayo, N. P. M., Nie, L., Peng, S., Islam, M. R., Sta. Cruz, P., Collard, B. C. Y., & Kato, Y. (2020). Identification and characterization of high- yielding, short-duration rice genotypes for tropical Asia. Crop Science, 60(5), 2241– 2250. 10.1002/csc2.20183

Wu, D., Liang, W., Zhu, W., Chen, M., Ferrándiz, C., Burton, R. A., Dreni, L., & Zhang, D. (2018). Loss of LOFSEP transcription factor function converts spikelet to leaf- like structures in rice. Plant Physiology, 176(2), 1646–1664. 10.1104/pp.17.00704

Xu, C., Liu, Y., Li, Y., Xu, X., Xu, C., Li, X., Xiao, J., & Zhang, Q. (2015). Differential expression of GS5 regulates grain size in rice. Journal of Experimental Botany, 66(9), 2611–2623. 10.1093/jxb/erv058

Yamaguchi, S., & Kamiya, Y. (2000). Gibberellin Biosynthesis: Its Regulation by Endogenous and Environmental Signals. In *Plant CellPhysiol* (Vol. 41, Issue 3). https://academic.oup.com/pcp/article/41/3/251/1828786

Yan, W. H., Wang, P., Chen, H. X., Zhou, H. J., Li, Q. P., Wang, C. R., Ding, Z. H., Zhang, Y. S., Yu, S. Bin, Xing, Y. Z., & Zhang, Q. F. (2011). A major QTL, Ghd8, plays pleiotropic roles in regulating grain productivity, plant height, and heading date in rice. Molecular Plant, 4(2), 319–330. 10.1093/mp/ssq070

Yang, Z., Sun, L., Zhang, P., Zhang, Y., Yu, P., Liu, L., Abbas, A., Xiang, X., Wu, W., Zhan, X., Cao, L., & Cheng, S. (2019). TDR INTERACTING PROTEIN 3, encoding a PHD-finger transcription factor, regulates Ubisch bodies and pollen wall formation in rice. Plant Journal, 99(5), 844–861. 10.1111/tpj.14365

Yano, M., Katayose, Y., Ashikari, M., Yamanouchi, U., Monna, L., Fuse, T., Baba, T., Yamamoto, K., Umehara, Y., Nagamura, Y., & Sasaki, T. (2000). Hd1 , a Major Photoperiod Sensitivity Quantitative Trait Locus in Rice, Is Closely Related to the Arabidopsis Flowering Time Gene CONSTANS. In The Plant Cell (Vol. 12). http://genes.mit.edu/GENSCAN.html

Zhang, Z. H., Zhu, Y. J., Wang, S. L., Fan, Y. Y., & Zhuang, J. Y. (2019). Importance of the interaction between heading date genes Hd1 and Ghd7 for controlling yield traits in rice. International Journal of Molecular Sciences, 20(3). 10.3390/ijms20030516

