## Supplementary figures for "Omics-driven Identification of Candidate Genes and SNP markers in a Major QTL Controlling Early Heading in Rice"

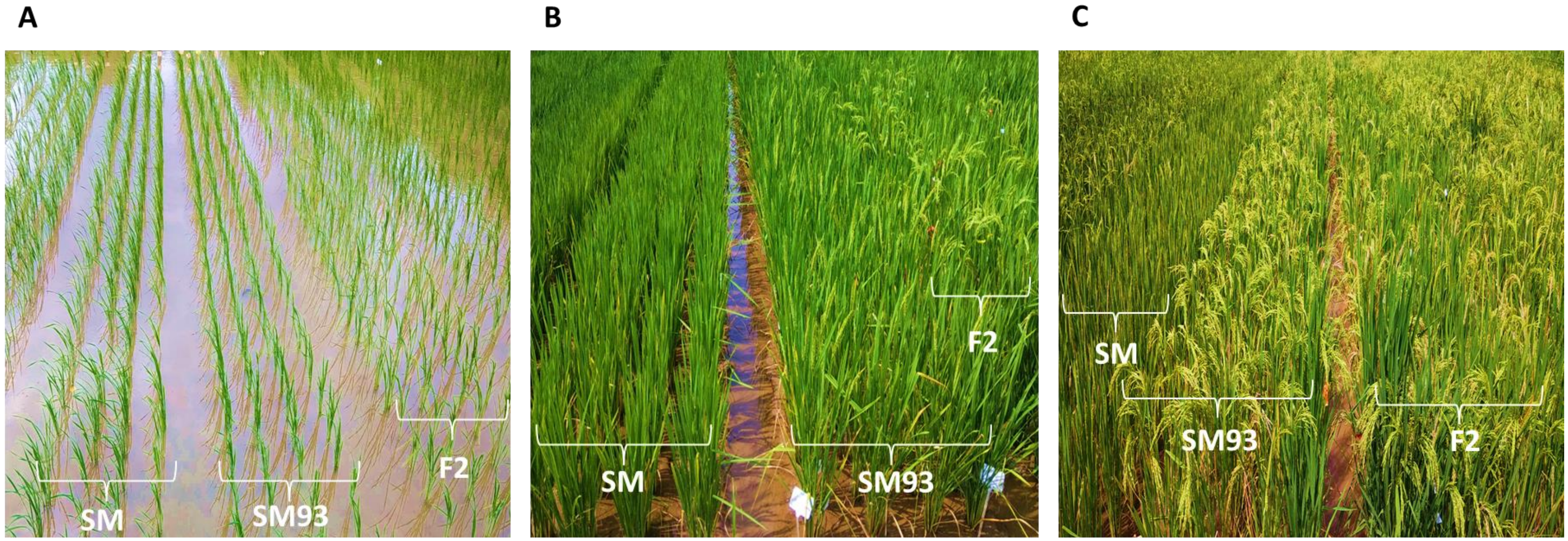


***Figure S1: Cultivation of SM, SM93 and F2 population in the field.*** ***(A)*** Plantlets in the vegetative stage, immediately after transplantation. ***(B)*** SM93 plants flower while the SM plants are still in the booting stage. The F2 generation displays heterogeneity in its flowering time. ***(C)*** All the SM93 plants with heavily loaded panicles, while the SM plants are in the initial stages of grain-filling. The F2 plants display variations in maturity.

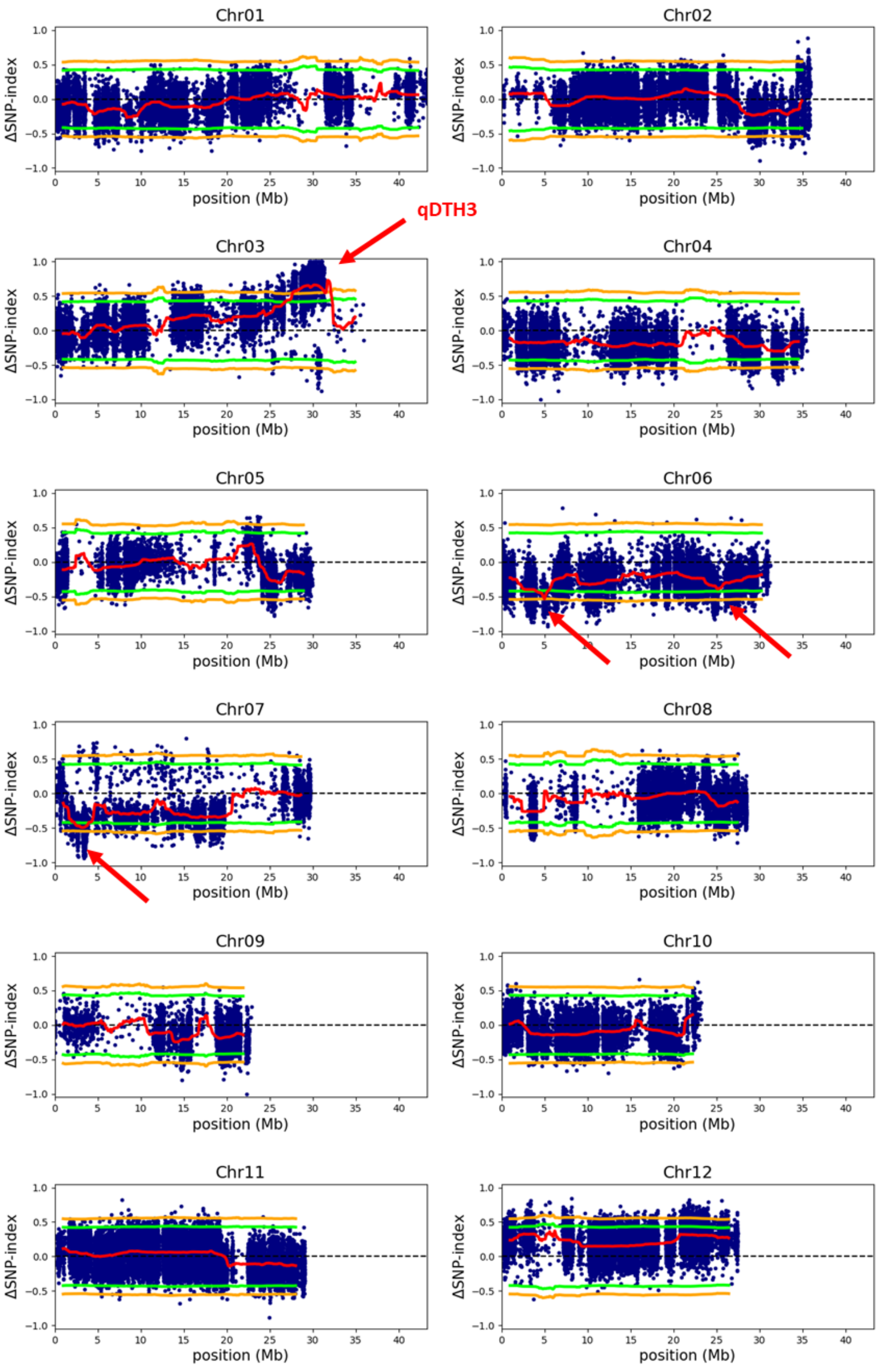


***Figure S2: QTL-seq Analysis of Kharif 2019's Population.*** Red arrows indicate detected QTLs.

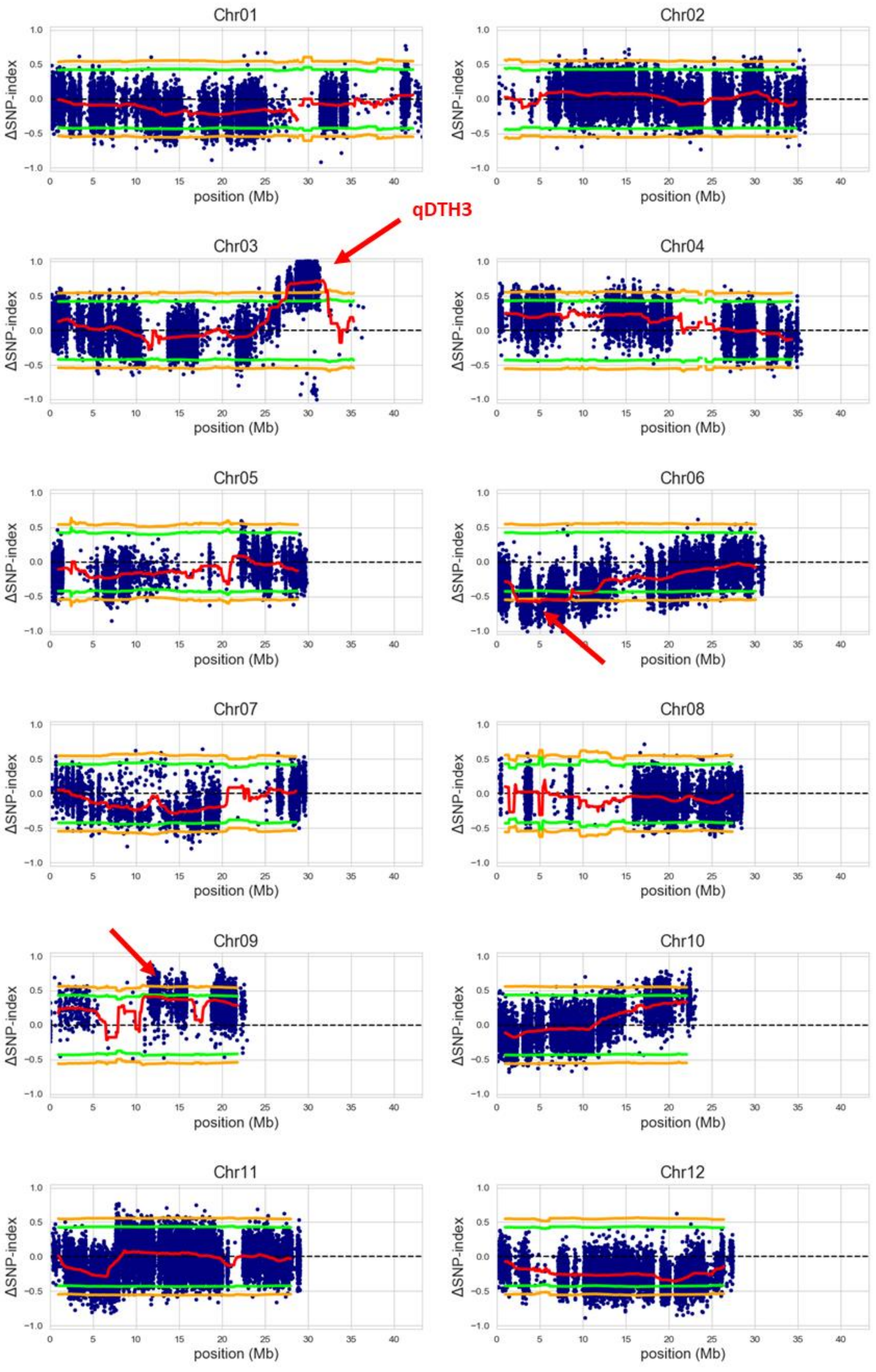


***Figure S3: QTL-seq Analysis of Kharif 2020's Population.*** Red arrows indicate detected QTLs.

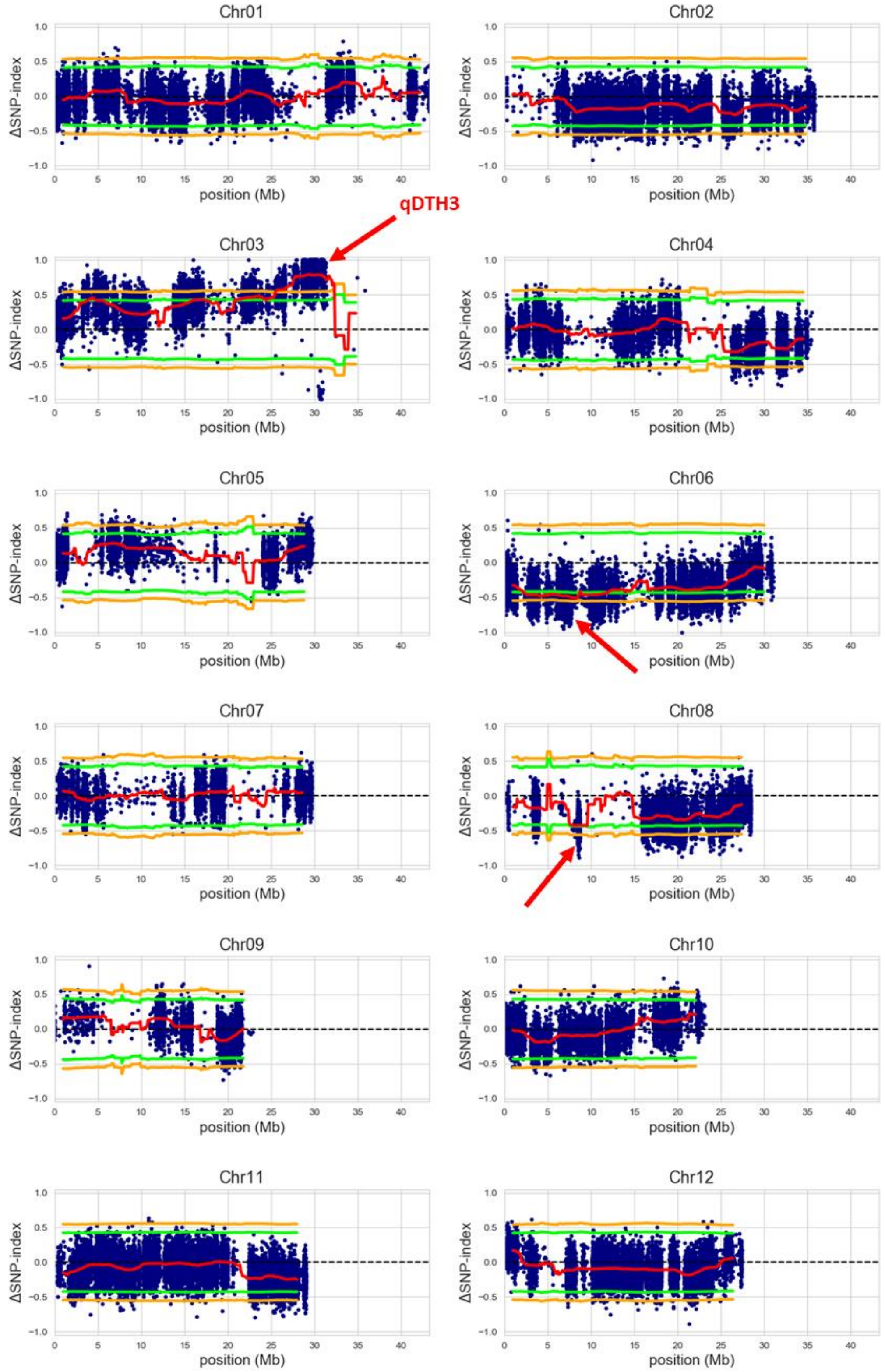


***Figure S4: QTL-seq Analysis of Kharif 2021's Population.*** Red arrows indicate detected QTLs.

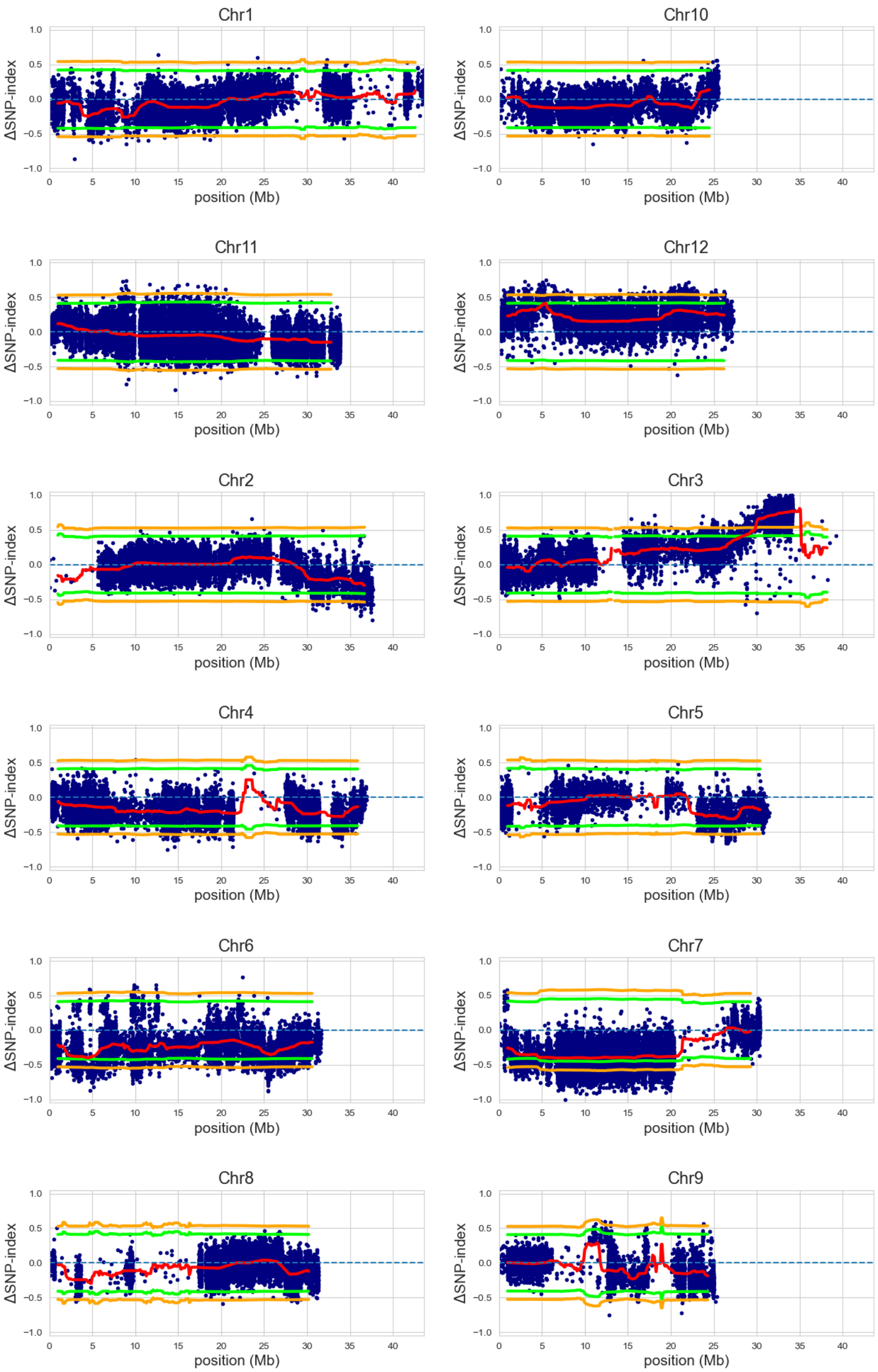


***Figure S5: QTL-seq Analysis of Kharif 2019's Population using the SM reference genome.***

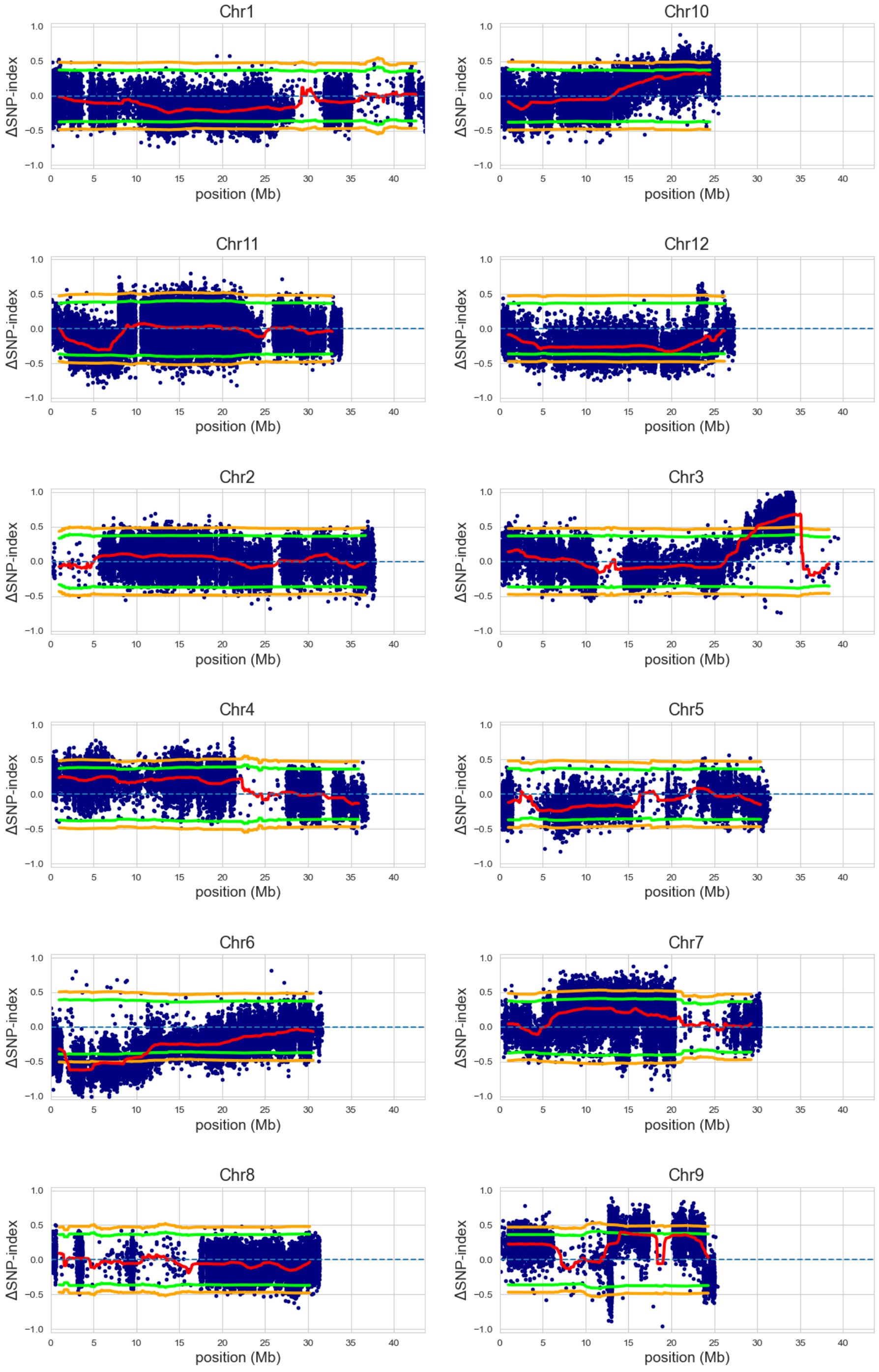


***Figure S6: QTL-seq Analysis of Kharif 2020's Population using the SM reference genome.***

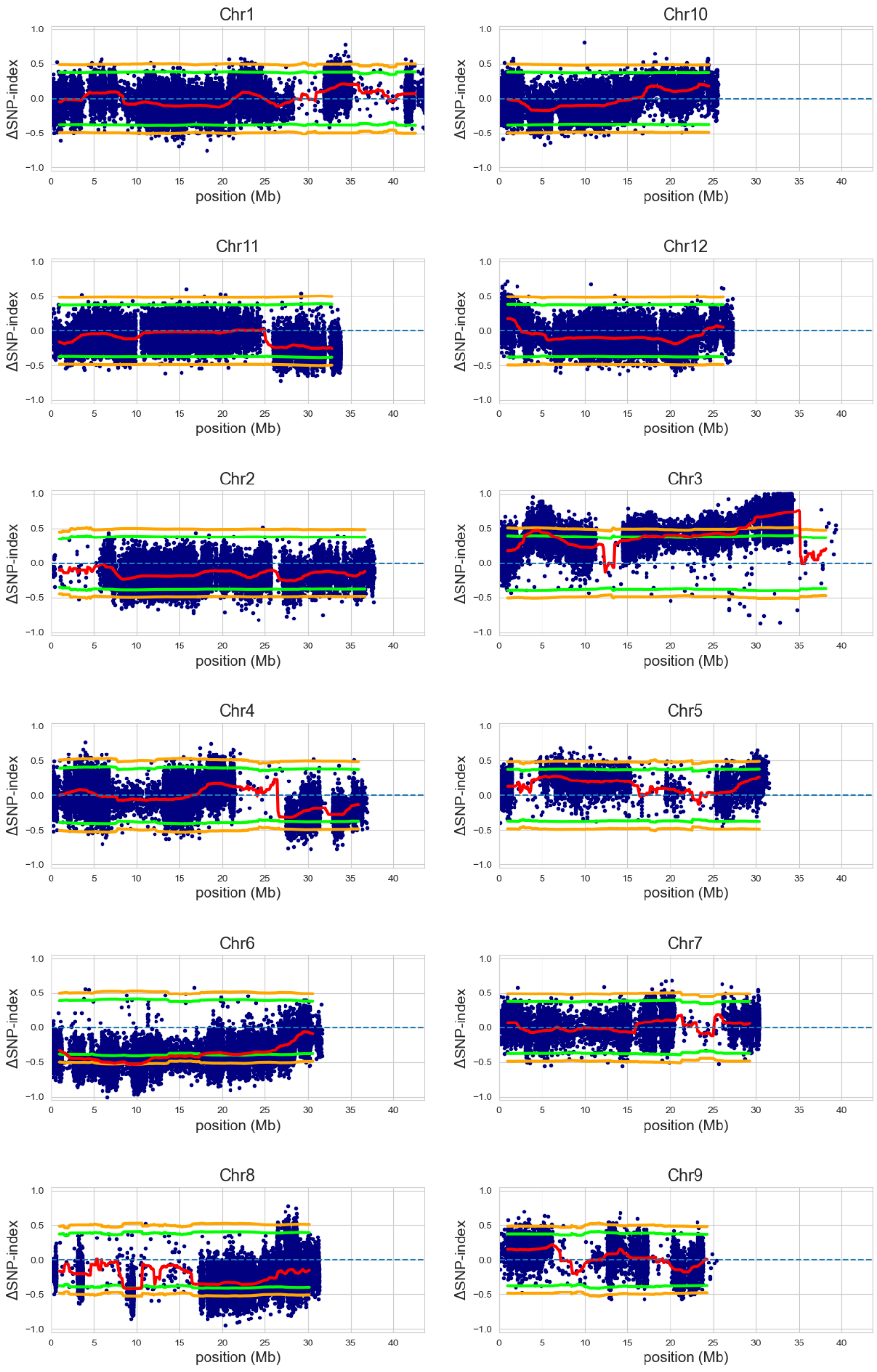


***Figure S7: QTL-seq Analysis of Kharif 2021's Population using the SM reference genome.***

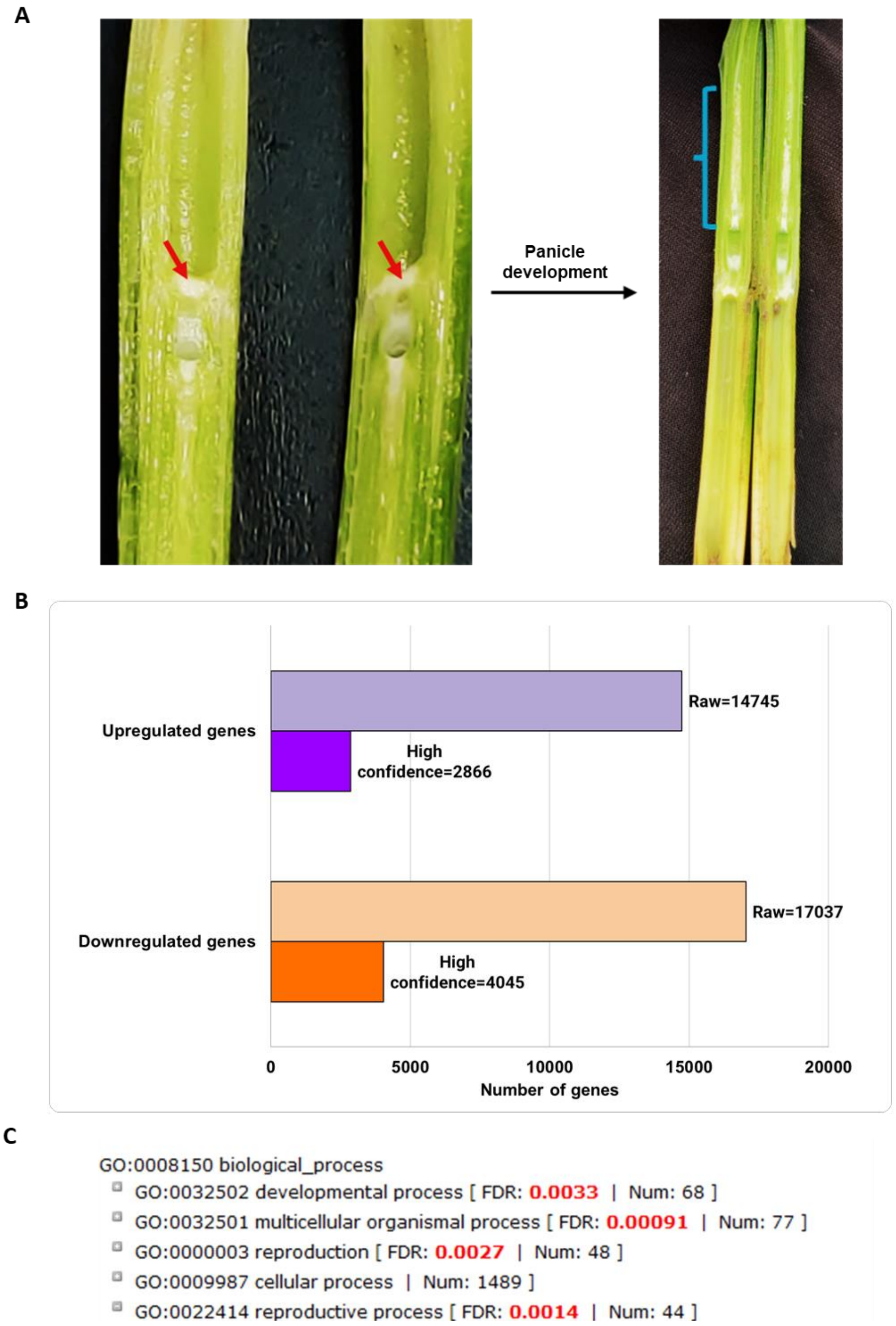


***Figure S8: Differential Gene Expression (SM93 vs SM) at the PI Stage.*** ***(A)*** Sampling of incipient panicle tissues. Red arrows indicate the incipient panicle, while the blue bracket indicates the growing panicle. ***(B)*** Global differences in gene expression between SM and SM93. ***(C)*** Gene Ontology Enrichment Analysis of DEGs.
